# KIF18A Maintains Kinetochore-Microtubule Attachments in CIN Cells by Limiting Microtubule Polymerization

**DOI:** 10.64898/2026.04.15.718686

**Authors:** Cindy Fonseca, Kira Fisher, Ethan Wagner, Sarah-Catherine Paschall, Haein Kim, Jason Stumpff

## Abstract

Chromosomal instability (CIN) generates vulnerabilities that can be therapeutically exploited, including sensitivity to inhibition of the kinesin motor KIF18A. However, the mechanistic basis for why a subset of CIN tumor cells depend on KIF18A remains unclear. Here, we compare mitotic phenotypes across KIF18A-sensitive and –insensitive cell models. In sensitive CIN cells, KIF18A inhibition leads to formation of polar chromosomes with unattached kinetochores, recruitment of spindle assembly checkpoint proteins, and prolonged mitotic arrest. Although KIF18A loss reduces kinetochore-microtubule stability in all cell lines, sensitive cells exhibit lower baseline attachment stability and heightened microtubule polymerization rates, predisposing them to attachment failure. Acute KIF18A inhibition disrupts maintenance of attachments after metaphase alignment, while reducing microtubule polymerization suppresses mitotic defects. These findings support a model in which CIN tumor cells rely on KIF18A to restrain excessive microtubule dynamics and maintain attachment.

## Introduction

Chromosome instability (CIN) and the mitotic errors that drive it are well-established contributors to tumor initiation, progression, and therapeutic resistance^1–3^. CIN not only fuels genetic heterogeneity but also exposes stressors that can be exploited therapeutically^4^. The kinesin-like motor protein KIF18A has emerged as a promising target for CIN-positive tumor cells^5–7^. Several small-molecule inhibitors of KIF18A have recently been developed and are advancing in clinical testing^8–12^. Nevertheless, only a subset of CIN tumor cells is sensitive to KIF18A loss of function, and the mechanistic basis of this selective vulnerability remains incompletely understood^5–7^.

KIF18A, a member of the kinesin-8 family, promotes chromosome alignment during mitosis by suppressing the dynamics of microtubules attached to kinetochores^13–15^. Loss of KIF18A in diploid somatic cells results in chromosome alignment defects and an increase in micronucleus formation, but mitosis proceeds with near normal timing^16^. In contrast, primordial germ cells and some CIN tumor cells exhibit prolonged mitotic arrest and reduced proliferation due to spindle assembly checkpoint activation^5–7,17,18^. The spindle assembly checkpoint inhibits the activity of the anaphase promoting complex (APC/C), an E3-ubiquitin ligase, to prevent cells from proceeding through the metaphase-to-anaphase transition until all kinetochores have established end-on attachments to spindle microtubules and are under tension^19–22^. Furthermore, co-inhibition of APC/C and KIF18A in CIN tumor cells or germ cells exacerbates the mitotic arrest phenotype^23,24^. These results suggest KIF18A may be required to establish or maintain attachments between kinetochores and microtubules in CIN cells and germ cells but not diploid somatic cells, however, the underlying basis for this selective requirement is not understood.

To address this question, we combined fixed– and live-cell imaging to examine mitotic spindle and chromosome dynamics across a panel of cell lines that exhibit varying levels of sensitivity to KIF18A inhibition. We found that when KIF18A is inhibited in sensitive CIN cells, kinetochore-microtubule attachments fail or cannot be sustained, producing polar chromosomes that recruit spindle assembly checkpoint proteins and induce prolonged mitotic arrest. These effects correlate with elevated levels of HEC1 phosphorylation, offloading of CENP-E from kinetochores, and increased microtubule polymerization rates. Furthermore, drug treatments that reduce microtubule polymerization also reduce the mitotic arrest caused by KIF18A inhibition. Taken together, these results support a model in which abnormally high microtubule polymerization rates in CIN cells renders them dependent on KIF18A to suppress dynamics near kinetochores and maintain attachments.

## Results

To uncover the basis of KIF18A sensitivity, we compared mitotic phenotypes between cell lines that display pronounced proliferation defects following KIF18A inhibition (hereafter, “sensitive”) and those that do not (“insensitive”). Sensitive lines included HeLa, MDA-MB-231, HT29, SKOV3, and OVCAR3, all previously shown to exhibit mitotic delays of varying severity upon KIF18A loss ^5,10,23^. Insensitive models included RPE-1, HCT116, and the CIN-positive colorectal cancer line SW480 ^5,10,23^. To test the effects of KIF18A loss of function, cell lines were treated with KIF18A specific or scrambled control siRNAs that have been previously validated ^5,10,23^. All cell lines exhibited the expected chromosome alignment defects after KIF18A knockdown (KD). In addition, we engineered HeLa and RPE-1 cells that express an siRNA-resistant GFP-KIF18A construct under a doxycycline-inducible promoter to analyze the ability of KIF18A expression to rescue any observed phenotypes^10,25^.

### CIN cells sensitive to KIF18A inhibition form polar chromosomes

Comparison of sensitive and insensitive mitotic cells after KIF18A KD revealed that, beyond alignment defects, sensitive lines displayed a high frequency of chromosomes positioned abnormally near the spindle poles (Figure 1A-C). Although occasional polar chromosomes were observed in insensitive lines, their frequency did not increase significantly after KIF18A KD (Figure 1B-C). Prior work indicates that these polar chromosomes often harbor kinetochore-microtubule attachment defects and recruit spindle assembly checkpoint (SAC) proteins, potentially explaining the prolonged mitotic arrest seen in sensitive CIN cells^26^.

**Figure 1.**
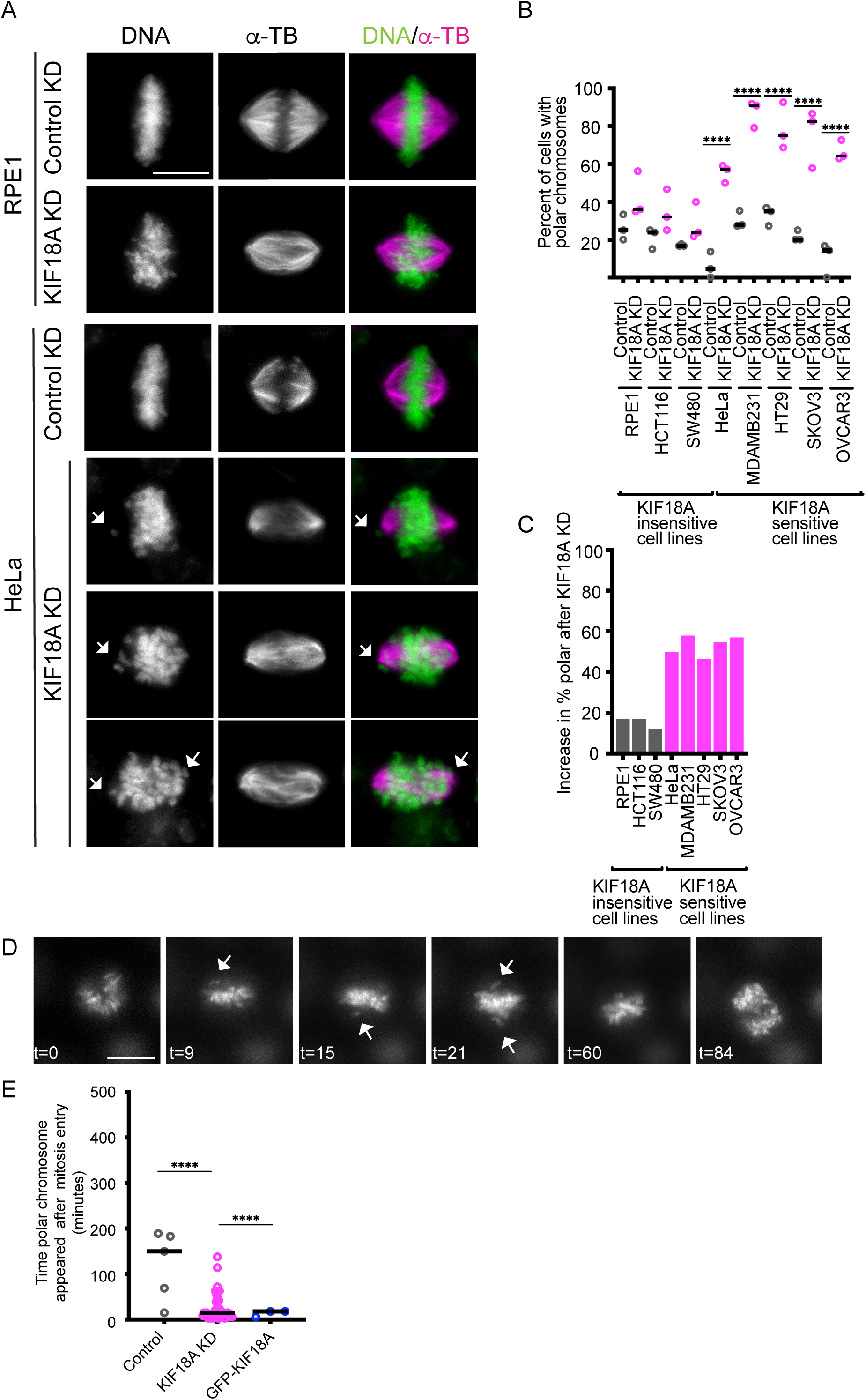
CIN cells sensitive to KIF18A inhibition form polar chromosomes. (A) Representative images of KIF18A-sensitive and –insensitive cells treated with control or KIF18A siRNAs for 24 hours before fixation. Microtubules were stained with anti-alpha-tubulin (αTb) antibodies and DNA was stained with DAPI. (B) Plot of the percentage of mitotic cells with polar chromosomes for each condition indicated. (C) Plot of fold-change in polar chromosomes for each cell type following KIF18A inhibition. (B-C) Data are from three biological replicates and the following number of cells for each condition: 86 (RPE1 control), 102 (RPE1 KIF18A KD), 102 (HCT116 control), 145 (HCT116 KIF18A KD), 70 (SW480 control), 127 (SW480 KIF18A KD), 112 (HeLa control), 184 (HeLa KIF18A KD), 74 (MDA-MB-231 control), 118 (MDA-MB-231 KIF18A KD), 100 (HT29 control), 230 (HT29 KIF18A KD), 14 (SKOV3 control), 57 (SKOV3 KIF18A KD), 20 (OVCAR3 control), 74 (OVCAR3 KIF18A KD). Statistical comparisons were made using a one-way ANOVA with Tukey’s multiple comparison. (D) Representative frames from live imaging of HeLa cells labeled with SPY-DNA. Time (t) in minutes is shown relative to mitotic entry. Arrows indicate polar chromosomes. (E) Plot of time after mitotic entry that polar chromosomes appeared in HeLa cells following the indicated treatments. Data are from three biological replicates and the following number of cells: 81 (control KD), 48 (KIF18A KD), and 44 (WT-GFP-KIF18A rescue in KIF18A KD). p-value style: <0.05 (*), <0.01 (**), <0.001 (***), and <0.0001 (****), not significant (>0.05) not shown. Scale bars = 10 μm

In HeLa cells, polar chromosomes were rare in control siRNA conditions but increased markedly to approximately 60% after KIF18A KD, identifying HeLa as a strong model for dissecting the origin of polar chromosomes (Figure 1B-C). Live-cell imaging of HeLa cells labeled with SPY-DNA revealed that polar chromosomes in KIF18A KD cells formed early in mitosis, whereas those observed in control cells appeared sporadically and without a temporal pattern (Figure 1D-E). Furthermore, induced expression of GFP-KIF18A in HeLa KIF18A KD cells significantly reduced the formation of polar chromosomes (Figure 1E). These observations indicate that loss of KIF18A drives an early defect in chromosome positioning, and polar chromosomes are not a secondary consequence of extended mitotic arrest.

### Polar chromosomes in KIF18A-depleted cells contribute directly to mitotic arrest

To investigate whether polar chromosome formation correlates with mitotic delay or arrest, we measured the time from DNA condensation to anaphase onset in live HeLa and RPE-1 KIF18A KD cells labeled with SPY-DNA. Consistent with previous work^17^, we found that KIF18A KD caused a significant mitotic delay in HeLa cells, which was rescued by induced expression of a GFP-KIF18A transgene (Figure 2A-B). In HeLa cells that completed mitosis during the filming period, the average time from DNA condensation to anaphase was 33.6 ± 1.6 min in control KD cells, 79.8 ± 10.0 min in KIF18A KD cells, and 36.6 ± 3.3 min in KIF18A KD cells expressing GFP-KIF18A. In contrast, KIF18A KD did not significantly alter mitotic timing in RPE-1 cells (Figure S1A-B).

**Figure 2.**
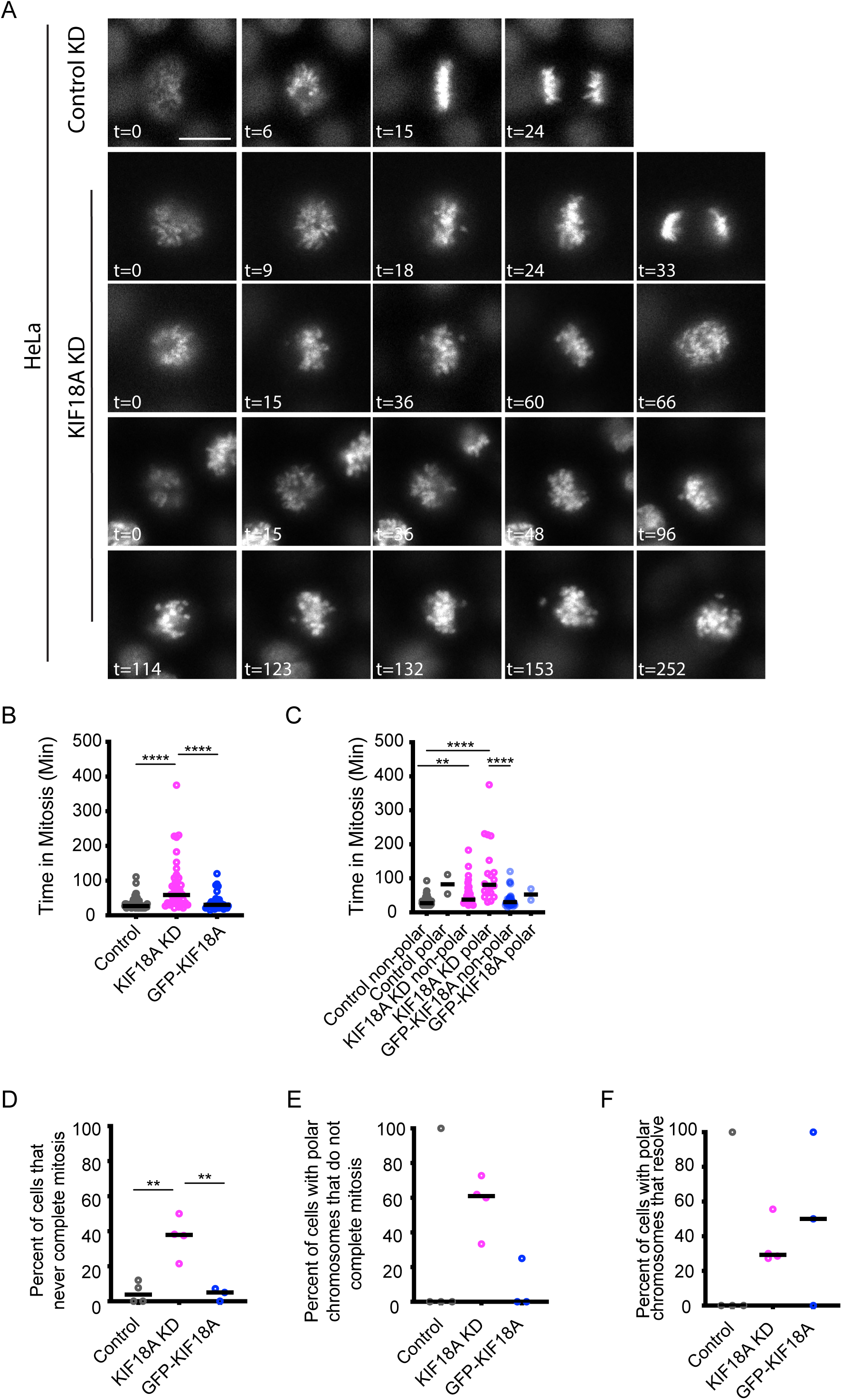
KIF18A-depleted cells with polar chromosomes exhibit prolonged mitotic arrest. (A) Representative still frames from time-lapse imaging of chromosomes in HeLa cells labeled with SPY-DNA. Time (t) in minutes relative to chromosome condensation is indicated. Scale bar = 10 μm. (B) Plots of time in mitosis, measured from chromosome condensation to anaphase onset, for HeLa cells treated with control siRNA, KIF18A siRNA, or KIF18A siRNA + induced expression of GFP– KIF18A. (C) Plot of time in mitosis for the same cells in (B) as a function of polar chromosome presence. (D-E) Plots of the percentage of all cells (D) or cells with polar chromosomes (E) that fail to complete mitosis during the imaging time and were in mitosis for at least 2 hours. (F) Plot of the percentage of cells from the indicated conditions that resolve polar chromosomes to the metaphase plate prior to anaphase. Data points in B-C represent individual cells. Data points in D-F represent independent experiments. All data are from 3 (GFP-KIF18A) or 4 (control KD and KIF18A KD) biological replicates. 81 cells (control KD), 48 cells (KIF18A KD), and 44 cells (KIF18A KD + GFP-KIF18A) were analyzed. Statistical comparisons were made using a Kruskal-Wallis Test with Dunn’s multiple comparison (B-C) or a one-way ANOVA with Tukey’s multiple comparison (D-F). p-value style: <0.05 (*), <0.01 (**), <0.001 (***), and <0.0001 (****), not significant (>0.05) not shown.

To assess a correlation between polar chromosomes and mitotic delay, we compared mitotic progression times in cells with or without polar chromosomes. Within the HeLa KIF18A KD population, cells that formed polar chromosomes and completed division remained in mitosis longer (114.4 ± 19.7 min) than those without polar chromosomes (55.2 ± 7.1 min) (Figure 2C). In contrast, no such trend was observed in RPE-1 cells (Figure S1A-B). We also observed that a notable fraction of HeLa cells were in mitosis for at least 2 hours but did not complete division during the imaging window, and the majority of these cells contained visible polar chromosomes (Figure 2D-E). However, a small fraction of HeLa KIF18A KD cells were able to resolve polar chromosomes by aligning them (Figure 2F). Together, these results support a strong association between KIF18A-dependent polar chromosome formation and prolonged mitotic arrest in HeLa cells.

Polar chromosomes frequently contain unattached kinetochores that recruit spindle assembly checkpoint (SAC) proteins. Consistent with this, KIF18A KD significantly increased the number of MAD1-positive kinetochores in HeLa cells, and this effect was fully rescued by expression of GFP-KIF18A (Figure S2A-B). Furthermore, the majority of MAD1-positive kinetochores were positioned off the metaphase plate, near the spindle poles (mean distance from the metaphase plate = 3.8±0.09 μm), confirming that polar chromosomes remain unattached and activate SAC signaling (Figure S2C). These observations demonstrate that chromosomes positioned near spindle poles in KIF18A-depleted cells exhibit attachment defects that contribute to prolonged mitotic arrest. In contrast, KIF18A KD did not significantly increase the number of MAD1-positive kinetochores in RPE-1 cells (Figure S2D).

### KIF18A inhibition decreases kinetochore-microtubule attachment

We next assessed the effects of KIF18A KD on kinetochore-microtubule attachments to determine if the unattached polar chromosomes occur due to a global decrease in attachments or if attachments are specifically affected on a subset of chromosomes in sensitive cells. To quantify stable kinetochore-microtubules, we treated HeLa and RPE1 cells with control or KIF18A siRNAs followed by CaCl_2_ treatment to destabilize non-kinetochore microtubules^27,28^. Cells were then fixed and stained with antibodies against α-tubulin. In both HeLa and RPE-1 cells, KIF18A knockdown (KD) reduced the abundance of CaCl_2_-resistant microtubules, measured either by total remaining tubulin or by plus-end fluorescence near kinetochores (Figure 3A-E). Expression of GFP-KIF18A restored microtubule stability in both cell models (Figure 3A-E). These results indicate that KIF18A is required for stable kinetochore-microtubule attachments in both sensitive and insensitive cells, which raises the question of why attachment defects only occur frequently after KIF18A KD in sensitive cells.

**Figure 3.**
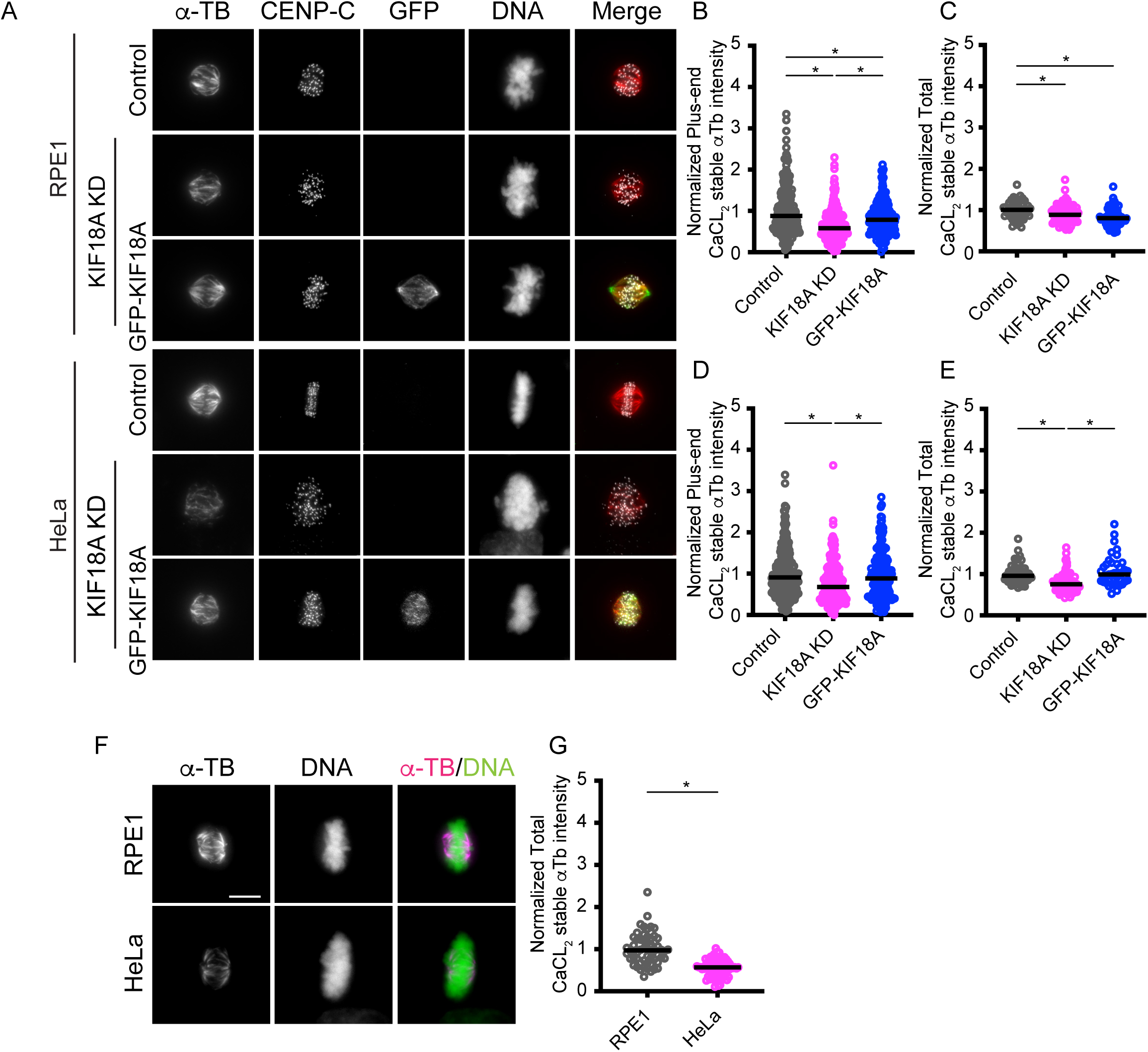
KIF18A inhibition decreases kinetochore-microtubule attachment. Representative RPE1 and HeLa cells fixed after CaCl_2_ treatment and stained for microtubules (α-TB), kinetochores (CENP-C), GFP-KIF18A (GFP), and DNA. (B-E) Plots of tubulin intensity following CaCl_2_ treatment for RPE1 (B-C) and HeLa cells (D-E) measured using an ROI near kinetochores (B, D,) or in the entire spindle (C, E). Data points represent individual measurements. Data are from three biological replicates and 52 cells (control KD, RPE1), 73 cells (KIF18A KD, RPE1), 70 cells (WT-GFP-KIF18A rescue in KIF18A KD, RPE1), 59 cells (control KD, HeLa), 77 cells (KIF18A KD, HeLa), and 41 cells (WT-GFP-KIF18A rescue in KIF18A KD, HeLa). (F) Representative images of RPE1 and HeLa cells following cold treatment. (G) Plot of tubulin fluorescence intensity measured in the spindle after CaCl_2_ treatment for each cell type. Data are from three biological replicates and 58 cells (RPE1) and 81 cells (HeLa). Statistical comparisons were made using a two-tailed t-test. p-value style: <0.05 (*), <0.01 (**), <0.001 (***), and <0.0001 (****), not significant (>0.05) not shown. Scale bars = 10 μm.

One possible explanation for this difference is that baseline kinetochore-microtubule stability is different in sensitive and insensitive cells. To compare the stable microtubules in HeLa and RPE-1 cells, we subjected otherwise untreated cells to CaCl_2_-treatment in parallel and measured α-tubulin fluorescence. The results revealed that RPE-1 cells contained a significantly higher amount of CaCl_2_-resistant microtubules than HeLa cells (Figure 3F-G). Thus, despite similar reductions in stability following KIF18A KD, this difference in baseline stability could explain why HeLa, but not RPE-1 cells, develop attachment defects and polar chromosomes when KIF18A is inhibited.

### KIF18A is required to maintain kinetochore-microtubule attachments

Prior studies of KIF18A have shown that the motor accumulates at the plus-ends of kinetochore-microtubules after attachment^14,17^, raising the question of whether KIF18A is required for initial attachment or maintenance of kinetochore-microtubule attachments. To address this question, we imaged metaphase-arrested HeLa cells treated with MG132 to generate aligned chromosomes with stable microtubule attachments and then acutely inhibited KIF18A using the small molecule sovilnesib^10^. These experiments were performed in HeLa cells expressing GFP-KIF18A, which can relocalizes to spindle poles after inhibition by sovilnesib^10^. This relocalization provides a visual readout of KIF18A inhibition in live cells. Following sovilnesib treatment, chromosomes moved away from the metaphase plate toward the spindle poles in a significantly higher fraction of cells compared with DMSO controls (Figure 4A-B). This behavior indicates that KIF18A activity is required for maintaining robust kinetochore-microtubule attachments, even after metaphase alignment has been achieved.

**Figure 4.**
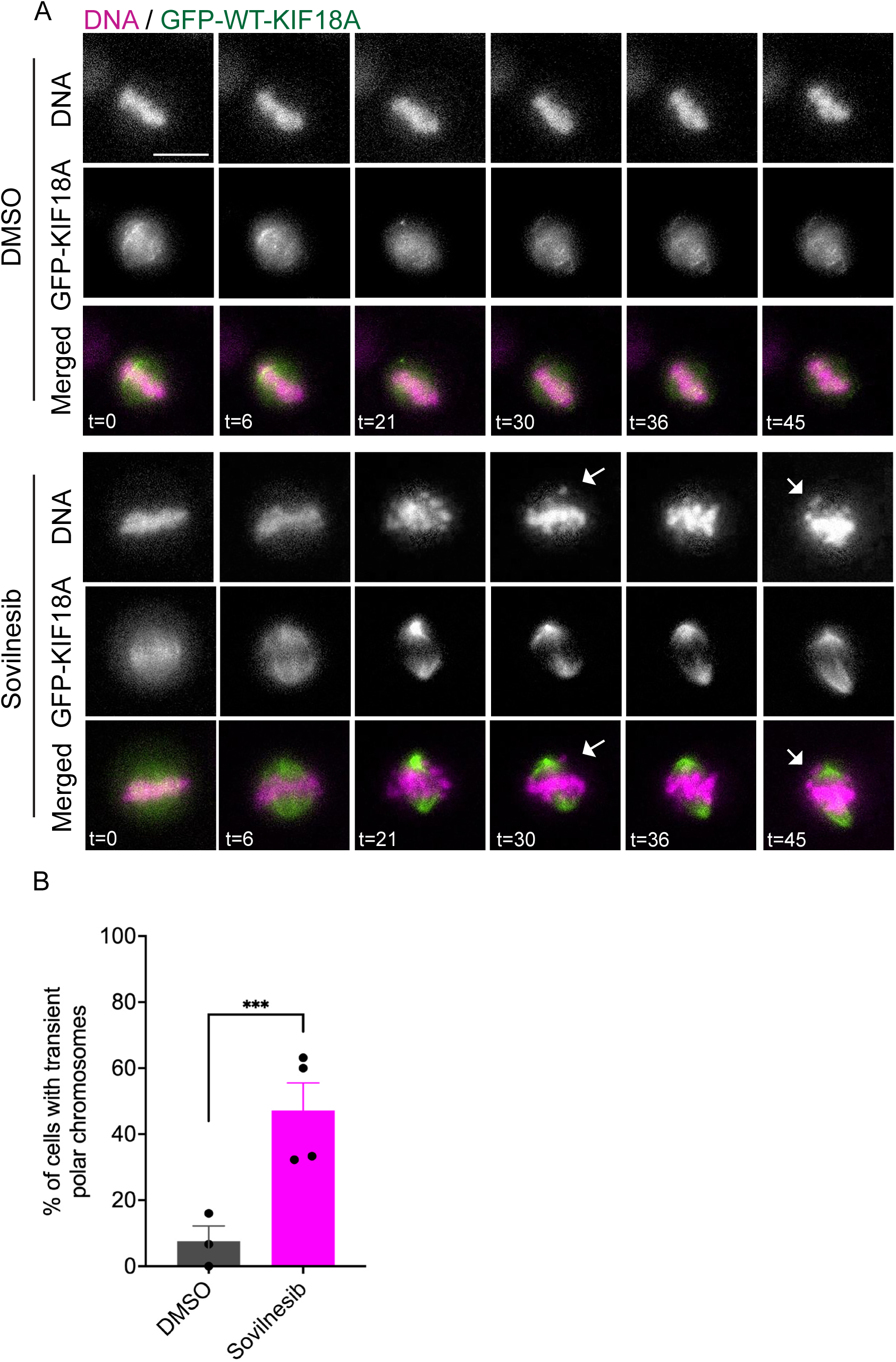
KIF18A is required to maintain established kinetochore-microtubule attachments. (A) Representative frames from time-lapse images of MG132-arrested HeLa cells expressing GFP-KIF18A and labeled with SPY-DNA following treatment with DMSO or the KIF18A inhibitor sovilnesib. Time (t) in minutes is measured relative to DMSO or sovilnesib addition. Arrows indicate polar chromosomes. Scale bar = 10 μm. (B) Plot of percentage of metaphase cells that form polar chromosomes following the indicated treatment. Mean ± SEM is shown with data points representing means from individual experiments. Data are from 3 (DMSO) or 4 (sovilnesib) biological replicates and 23 cells (DMSO) and 68 cells (sovilnesib). Statistical comparisons were made using a Welch’s t-test. p-value style: <0.05 (*), <0.01 (**), <0.001 (***), and <0.0001 (****), not significant (>0.05) not shown.

### HEC1 phosphorylation remains at prometaphase levels following KIF18A inhibition

To determine the molecular basis for kinetochore-microtubule attachment defects in KIF18A inhibited cells, we first investigated effects on the HEC1 complex. The HEC1 complex mediates microtubule binding at kinetochores, and its affinity for microtubules is regulated by phosphorylation of several residues in the N-terminus, with increased phosphorylation leading to reduced affinity^29–31^. During mitotic progression, HEC1 phosphorylation is reduced as stable kinetochore-microtubule attachments increase^32^. To determine whether KIF18A affects phospho-regulation of HEC1, we measured immunofluorescent signal from antibodies that recognize phosphorylated Ser44 or Ser55. In metaphase-arrested HeLa cells, KIF18A knockdown (KD) increased phospho-HEC1 to levels comparable to those observed in prometaphase cells, and expression of GFP-KIF18A reduced this phosphorylation, restoring it to metaphase levels (Figure 5A-C). These results indicate KIF18A is required for timely HEC1 dephosphorylation during metaphase, which likely contributes to reduced kinetochore-microtubule attachment stability.

**Figure 5.**
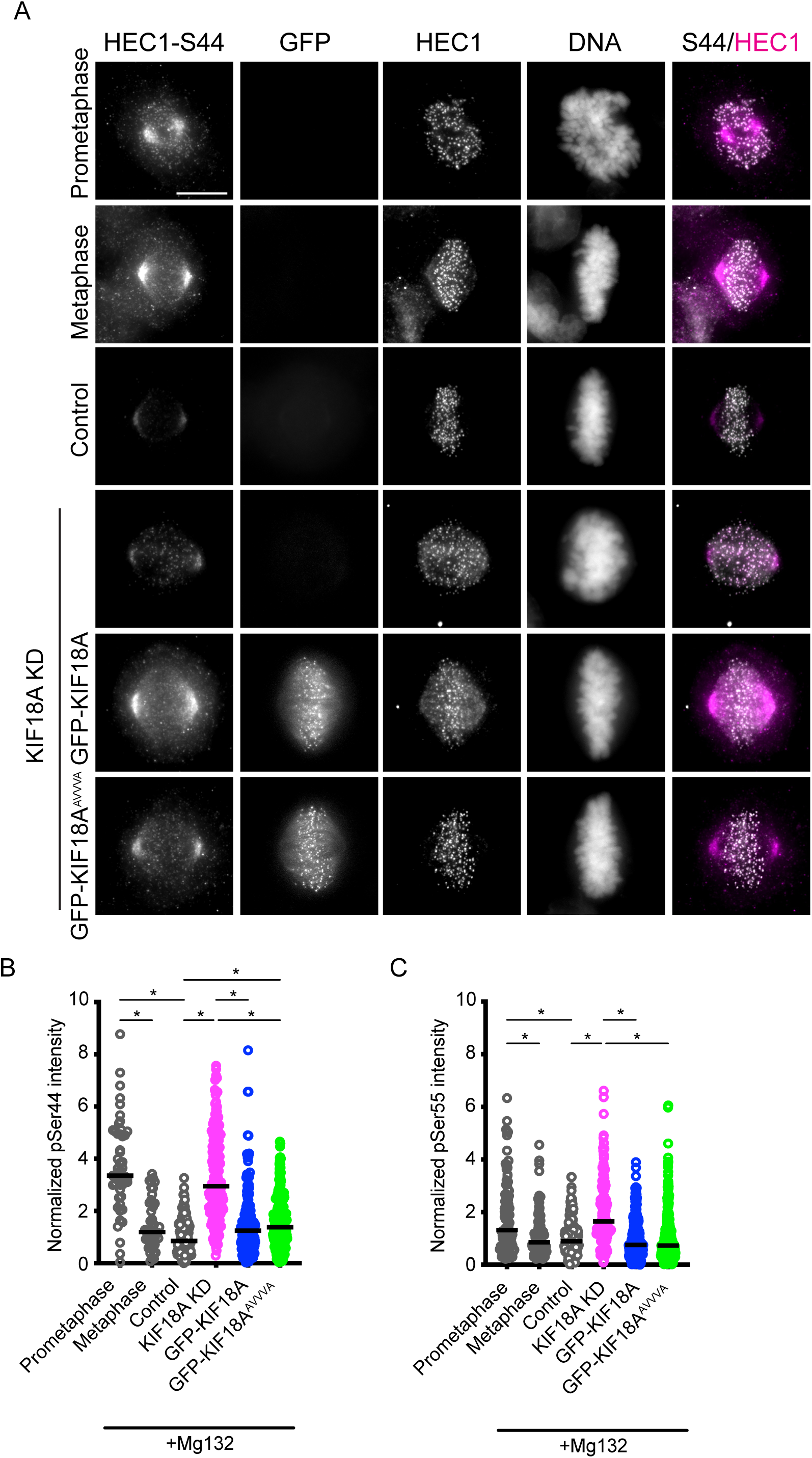
HEC1 phosphorylation remains at prometaphase levels following KIF18A inhibition. (A) Representative images of asynchronously dividing HeLa cells in the indicated mitotic stages or MG132-arrested cells treated with control or KIF18A siRNAs and induced to express WT GFP-KIF18A or a GFP-KIF18A containing mutations in the PP1-binding domain (GFP-KIF18A^AVVVA^). Cells were fixed and stained with antibodies against HEC1, phosphorylated serine 44 of HEC1 (HEC1-S44), and GFP. DNA was labeled with DAPI. Scale bar = 10 μm. (B-C) Plots showing fluorescence intensities measured for phosphorylated HEC1 S44 (B) and S55 (C) as a ratio to total HEC1 signal from the same ROI. Data points represent kinetochore measurements. Data are from three biological replicates and 51 (S44) and 65 (S55) kinetochores (prometaphase); 47 (S44) and 64 (S55) kinetochores (metaphase); 166 (S44) and 137 (S55) kinetochores (control KD); 203 (S44) and 135 (S55) kinetochores (KIF18A KD); 165 (S44) and 152 (S55) kinetochores (KIF18A KD, GFP-KIF18A); and 235 (S44) and 303 (S55) kinetochores (KIF18A KD, GFP-KIF18A^AVVVA^). Statistical comparisons were made using a one-way ANOVA with Tukey’s multiple comparison. p-value style: <0.05 (*), <0.01 (**), <0.001 (***), and <0.0001 (****), not significant (>0.05) not shown.

HEC1 is dephosphorylated through the activity of protein phosphatase 1 (PP1), and KIF18A has an established PP1-binding site in its tail domain^33^. To assess whether direct binding of PP1 to KIF18A is required for its ability to promote HEC1 dephosphorylation, we analyzed phospho-HEC1 levels in cells expressing a PP1-binding-deficient KIF18A mutant, GFP-KIF18A^AVVVA 33,34^. We observed that GFP-KIF18A^AVVVA^ reduced phospho-HEC1 to similar levels as wild type GFP-KIF18A, indicating that PP1 recruitment to KIF18A is not required for the motor’s role in moderating HEC1 phosphorylation (Figure 5A-C). Together, these findings suggest that attachment defects in KIF18A-depleted cells correlate with persistence of prometaphase-like HEC1 phosphorylation states, but that KIF18A likely does not directly regulate HEC1 phosphorylation.

### CENP-E is prematurely offloaded from kinetochores in KIF18A-depleted cells

Several studies have implicated a functional relationship between KIF18A and the kinesin-7 motor CENP-E during mitosis^35–37^. CENP-E localizes to kinetochores, promotes end-on attachments between kinetochores and spindle microtubules, and facilitates congression of unattached kinetochores to the metaphase plate^38–41^. Thus, loss of CENP-E activity could contribute to the decreased attachment and polar chromosome phenotypes observed in KIF18A KD cells. In asynchronously dividing cells, CENP-E levels at kinetochores were reduced following KIF18A knockdown (KD) in HeLa but not RPE1 cells (Figure 6A-C). Expression of GFP-KIF18A in HeLa cells treated with KIF18A siRNAs also restored, and in some cases increased, CENP-E levels at kinetochores (Figure S3A-B). We also found that CENP-E levels were reduced at all kinetochores, and levels were similar on kinetochores near the spindle poles or near the metaphase plate (Figure 6B-C).

**Figure 6.**
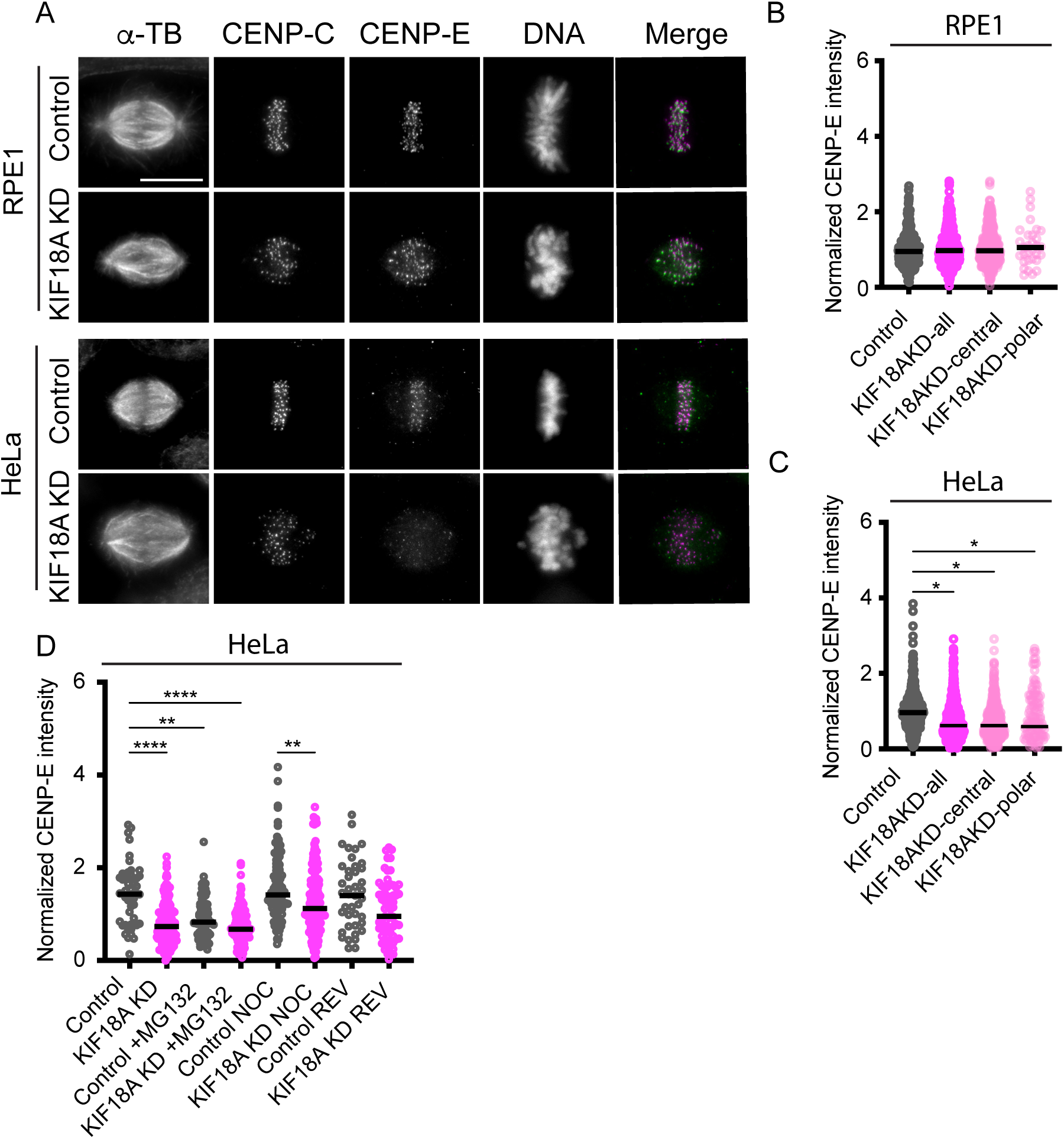
CENP-E is prematurely offloaded from kinetochores in KIF18A-depleted cells. (A) Representative images of control and KIF18A siRNA treated RPE1 and HeLa cells that were fixed and labeled with antibodies to alpha-tubulin (αTb), CENP-C, and CENP-E. DNA is labeled with DAPI. Scale bar = 10 μm. (B-C) Plots of CENP-E fluorescence intensity at all kinetochores or those localized to near the spindle poles (polar) or metaphase plate (central) normalized to levels in control siRNA treated cells in RPE1 (B) and HeLa cells (C). Data points represent individual kinetochores and lines indicate median. Data are from three biological replicates and the following number of kinetochores: 426 (RPE1 control); 601 (RPE1 KIF18A KD all), 570 (RPE1 KIF18A KD central); 30 (RPE1 KIF18A KD polar), 480 (HeLa control); 684 (HeLa KIF18A KD all), 555 (HeLa KIF18A KD central); and 129 (HeLa KIF18A KD polar). (D) Plot of normalized CENP-E fluorescence intensity at kinetochores in HeLa cells treated with control or KIF18A siRNAs and MG132, nocodazole (NOC), or reversine (REV). Data points represent individual cells and lines indicate median. Data are from three biological replicates and the following number of cells: 56 (control); 172 (KIF18A KD), 79 (control + MG132); 156 (KIF18A KD + MG132), 109 (control + NOC); 171 (KIF18A KD + NOC); 43 (control + REV); and 69 (KIF18A KD + REV). Statistical comparisons for B-D were made using Kruskal-Wallis with Dunn’s multiple comparisons test. p-value style: <0.05 (*), <0.01 (**), <0.001 (***), and <0.0001 (****), not significant (>0.05) not shown.

CENP-E is normally recruited directly to kinetochores and then is removed via dynein-mediated transport along kinetochore-microtubules once stable attachments are established^42,43^. To determine whether CENP-E reduction at kinetochores in KIF18A KD HeLa cells reflects defective recruitment or premature removal of CENP-E, we examined how CENP-E levels were affected by drug treatments that alter mitotic progression or microtubule attachment status. Arresting cells in metaphase with stable attachments via MG132 treatment reduced CENP-E levels in control KD cells to levels that were comparable to those in KIF18A-depleted cells with or without MG132, indicating mitotic arrest results in reduced CENP-E at kinetochores (Figure 6D and Figure S3C). On the other hand, treatment with the MPS1 inhibitor reversine, which alleviates spindle assembly checkpoint-mediated arrest, increased CENP-E levels at kinetochores in KIF18A KD HeLa cells (Figure 6D and Figure S3C). To evaluate whether microtubule attachment status influences CENP-E retention, we treated cells with nocodazole to depolymerize microtubules and inhibit CENP-E offloading. CENP-E levels in nocodazole-treated KIF18A KD cells were comparable to those in control KD, nocodazole-treated cells, suggesting that initial recruitment of CENP-E remains intact in the absence of KIF18A activity (Figure 6D and Figure S3C). Together, these results indicate that reduced CENP-E at kinetochores in KIF18A-depleted HeLa cells primarily reflects premature offloading driven by extended mitotic arrest, rather than a defect in recruitment. Importantly, this suggests that decreased CENP-E is not the primary cause of polar chromosome formation during early mitosis in KIF18A-inhibited cells.

### Increased microtubule polymerization following KIF18A inhibition contributes to attachment defects

Chromosomally unstable (CIN) tumor cells exhibit elevated microtubule polymerization rates, which contribute to chromosome instability^44^. KIF18A functions to suppress microtubule dynamics, including the reduction of plus-end polymerization^13,15,45^. We therefore tested whether inhibition of KIF18A with sovilnesib further increases polymerization in CIN-positive cells. Acute inhibition with sovilnesib increased EB3-labeled microtubule plus-end growth rates in CIN cells with significant increases measured in the most KIF18A-sensitive lines (HT29, OVCAR3, and MDA-MB-231) (Figure 7A). In contrast, inhibition of KIF18A did not increase microtubule polymerization in RPE1 cells (Figure 7A).

**Figure 7.**
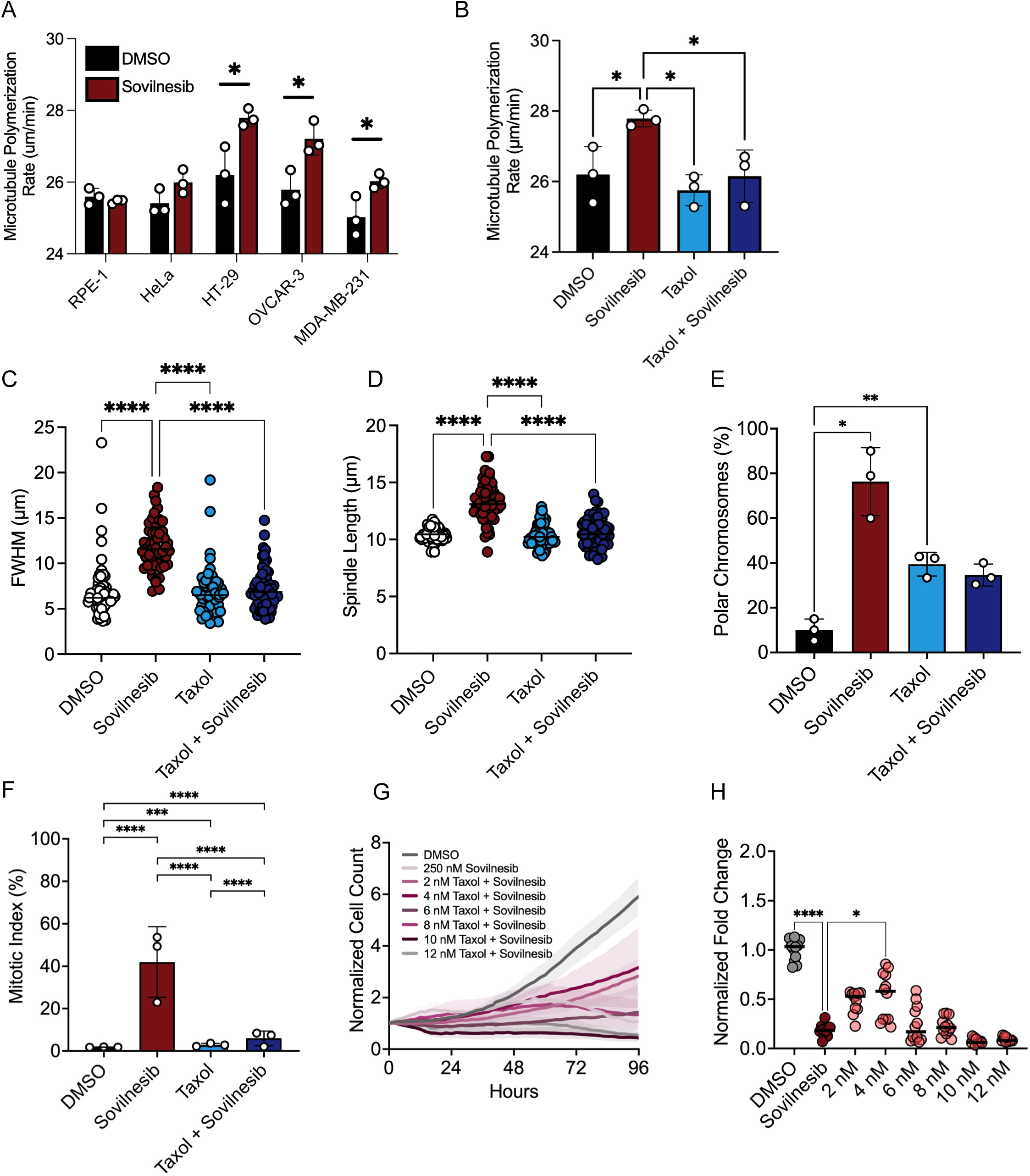
Increased microtubule polymerization following KIF18A inhibition contributes to attachment defects. (A) Measurement of mitotic microtubule plus-end polymerization rates (μm/min) across the indicated CIN cell types after 24-hour treatment with DMSO (black) or 250 nM Sovilnesib (red). Each dot represents average polymerization rate from experimental replicates (n=11-23 cells from three independent experiments). Data were compared using a two-way ANOVA with Sidak’s multiple comparisons test. (B) Measurement of mitotic microtubule plus-end polymerization rates (μm/min) in HT-29 cells after 24-hour treatment with DMSO (black), 250 nM Sovilnesib (red), 2 nM Taxol (light blue), or Sovilnesib-Taxol combination (dark blue). Each dot represents average polymerization rate from experimental replicates (n=9-23 cells from three independent experiments). Data were compared using a one-way ANOVA with Tukey’s post hoc test. (C-D) Plots of full-width at half maximum (FWHM) of centromere fluorescence distribution along the length of the spindle (C) and spindle length (D) measured in HT-29 cells fixed and stained after 24-hours of the indicated treatments. Each dot represents a single cell (n=17-21 cells from three independent experiments). Mean ± standard deviation is displayed. Data were compared by Kruskal–Wallis with Dunn’s multiple comparisons test. (E-F) Plots showing percentage of mitotic cells with polar chromosomes (E) and mitotic index (% of total cells in mitosis) (F) in HT-29 cells after 24 hours of indicated treatment. Each dot represents the mean from a biological replicate. Bars are mean ± standard deviation. Data were compared using a chi-squared test. (G) Plot of normalized cell count over time for HT-29 cells treated with indicated concentrations of Sovilnesib and Taxol. Lines represent mean, and shaded areas represent standard deviation. (H) Plot of normalized HT-29 cell count (shown as a % of DMSO control) as a function of Sovilnesib and Taxol concentrations. Data are from three biological replicates. Each dot represents an individual well (9 wells per condition) and bars indicate mean values. Data were compared by Mann–Whitney test. p-value style: <0.05 (*), <0.01 (**), <0.001 (***), and <0.0001 (****), not significant (>0.05) not shown.

To determine if the increased microtubule polymerization rates induced by sovilnesib contribute to attachment defects, we treated HT29 cells with a low dose of taxol (2nM), which lowered microtubule polymerization rates in sovilnesib-treated cells back to a level comparable to rates measured in control cells (Figure 7B). Notably, this concentration of taxol did not significantly reduce proliferation of HT29 cells on its own (Figure S4A). However, we found that combining 2-4nM taxol with 250nM sovilnesib reduced the chromosome alignment, spindle length, polar chromosome, mitotic arrest, and proliferation defects caused by sovilnesib alone (Figure 7C-H). These findings support a model in which CIN tumor cells rely on KIF18A to moderate their inherently high microtubule polymerization rates, and loss of this activity leads to destabilized kinetochore-microtubule attachments.

## Discussion

Our findings support an integrated model in which elevated microtubule polymerization in chromosomally unstable (CIN) tumor cells creates a dependence on KIF18A to suppress polymerization and stabilize kinetochore-microtubule attachments. In KIF18A-sensitive cells, inhibition of the motor prevents the formation and maintenance of robust kinetochore-microtubule attachments, producing early, unattached polar chromosomes that recruit spindle assembly checkpoint components and induce prolonged mitotic arrest. In contrast, cells with intrinsically higher baseline attachment levels, such as RPE-1 cells, tolerate a similar decrease in stability without crossing the threshold that triggers checkpoint protein recruitment to a subset of kinetochores and subsequent mitotic arrest.

Several lines of evidence support that KIF18A is required for establishment and maintenance of robust kinetochore-microtubule attachments. KIF18A inhibition decreased CaCl_2_-resistant kinetochore-microtubules in both sensitive and insensitive cells, indicating that the motor plays a general role in promoting kinetochore-microtubule attachment. Furthermore, acute inhibition of KIF18A resulted in chromosomes detaching and moving off the metaphase plate in HeLa cells, demonstrating a continued dependence on KIF18A for maintaining attachments even after alignment. Reduced attachments after KIF18A depletion were also accompanied by elevated phospho-HEC1 levels at kinetochores, consistent with a reduced microtubule-affinity state reminiscent of prometaphase.

KIF18A does not localize near kinetochores in the absence of microtubules, and its PP1-binding domain does not significantly contribute to dephosphorylation of HEC1, suggesting that the motor does not directly link microtubules to kinetochores or directly regulate the affinity of kinetochores for microtubules^14,17,33^. How then does the motor promote kinetochore-microtubule attachment? Both in cells and in vitro, KIF18A accumulates at the plus-ends of microtubules and suppresses dynamics^13,15,45^. Loss of such an activity at kinetochore-microtubule plus-ends could explain previous reports of increased microtubule turnover at kinetochores and reduced presence of depolymerizing microtubules on the anti-poleward moving (trailing) kinetochores during metaphase chromosome oscillations in KIF18A-depleted cells^46,47^. This latter change would reduce drag forces during chromosome movements, explaining the faster kinetochore velocities and reduced tension across kinetochores observed in KIF18A-depleted cells^14,15^. Collectively, increased kinetochore-microtubule turnover and reduced tension in KIF18A inhibited cells are expected to result in fewer microtubule attachments per kinetochore and increased HEC1 phosphorylation by Aurora B kinase^30,31,48^. Thus, we propose that KIF18A’s regulation of kinetochore-microtubule dynamics is necessary for maintaining their attachment to kinetochores.

Combining this loss of kinetochore-microtubule dynamics regulation with the altered spindle microtubule dynamics exhibited by CIN cells can explain the selective KIF18A-dependency of CIN cells. CIN cells have been shown to exhibit faster than normal microtubule polymerization rates during mitosis^44^, and we found that KIF18A inhibition exacerbates this defect, which is predicted to elevate turnover of kinetochore-microtubule attachments. Notably, only a small subset of kinetochores in CIN cells localized near spindle poles and recruited the spindle checkpoint protein MAD1. This is consistent with a threshold of kinetochore-microtubule attachment being required to satisfy the checkpoint ^49^, and we propose that the combination of altered microtubule dynamics and KIF18A inhibition in CIN cells pushes some kinetochores below this threshold. In further support of this model, reducing microtubule polymerization with low-dose taxol in sensitive cells reduced the mitotic defects caused by KIF18A inhibition, leading to increased proliferation. Together, these data define a coherent mechanistic model linking polymerization dynamics, attachment stability, and mitotic arrest in KIF18A-sensitive cells.

This model is also consistent with previous work demonstrating that perturbing kinetochore-microtubule attachments or spindle assembly checkpoint function alters cellular responses to KIF18A inhibition. For example, increased microtubule polymerization and decreased kinetochore-microtubule attachment in KIF18A inhibited cells would explain sensitivity to changes in anaphase-promoting complex (APC/C) activity, as this would affect the duration of mitotic delay resulting from the primary attachment defects^23,24^. Similarly, loss of KIF18A’s kinetochore-microtubule stabilizing activity would be predicted to enhance effects of other mutations that reduce attachment, as recently shown for a hypomorphic CENP-C mutant and inhibition of CENP-E^37^. Our studies of CENP-E also provide further insight into its functional relationship with KIF18A, suggesting that the mitotic arrest caused by KIF18A loss in sensitive cells leads to CENP-E offloading from kinetochores. This, in turn, is predicted to exacerbate defects in resolving polar chromosomes, which is an established function for CENP-E.

Our proposed model also yields testable predictions and suggests biomarkers for stratifying tumors likely to respond to KIF18A inhibition. For example, baseline measures of kinetochore-microtubule stability, such as CaCl_2_-resistant tubulin intensity or EB3 comet velocities, may correlate with sensitivity. Elevated prometaphase-like phosphorylation states at kinetochores, such as increased phospho-HEC1, may also serve as mechanistic readouts of attachment instability. Furthermore, tumors with high baseline microtubule stability, or with regulatory mechanisms that buffer polymerization dynamics, may show reduced responses to KIF18A inhibitors.

This framework also suggests rational combinatorial treatment strategies for CIN tumors. Agents that temper microtubule polymerization, such as low-dose microtubule-stabilizing drugs, or those that reduce kinetochore phosphorylation, such as Aurora kinase inhibitors, may restore or maintain attachment stability sufficiently to alter the magnitude or duration of mitotic arrest. Conversely, combinations that further elevate polymerization or disrupt microtubule capture pathways, such as microtubule depolymerizing drugs or higher doses of microtubule-stabilizing compounds, might selectively exacerbate attachment failures in KIF18A-dependent tumors. Optimizing the timing and dosing of such agents will be necessary to enhance therapeutic windows while minimizing toxicity.

Future studies should prioritize identifying genomic or proteomic correlates of elevated polymerization. For example, tubulin isoform expression ratios, altered tubulin homeostasis pathways, or regulators such as Aurora A could potentially be used to identify sensitive tumor cells. Establishing quantitative biomarkers of kinetochore-microtubule stability in patient-derived models will be critical for translating these findings. Furthermore, therapeutic evaluations should test combination regimens that selectively push CIN cells beyond their tolerance for attachment turnover while sparing non-tumor cells. Together, these directions will help refine patient selection strategies and accelerate the development of KIF18A-targeted therapies.

## Resource Availability

### Lead contact

Jason Stumpff

### Materials availability

All unique/stable reagents generated in this study are available from the lead contact without restriction.

### Data and code availability

All data reported in this paper will be shared by the lead contact upon request.

## Acknowledgements

This work was supported by NIH R35 GM144133 to JS. We thank Jennifer DeLuca for providing phospho-specific antibodies to HEC1.

## Author Contributions

Conceptualization, CF, KF, EW, SP, HK, JS; formal analysis, CF, KF, EW, JS; funding acquisition, JS; investigation, CF, KF, EW, SP, HK; supervision, JS; visualization, CF, KF, EW; writing-original draft, CF, KF, EW, JS; writing-review & editing, CF, KF, EW, SP, HK, JS.

## Declaration of Interests

JS is the inventor on a patent related to this work: US20230233565A1.

## Supplemental information

Document S1. Figures S1-S4

## Figures

**Figure S1.** KIF18A KD does not induce polar chromosomes in RPE1 cells. (A) Representative still frames from time-lapse imaging of chromosomes in RPE1 cells labeled with SPY-DNA. Time (t) in minutes relative to chromosome condensation is indicated. Scale bar = 10 μm. (B) Plot of time in mitosis, measured from chromosome condensation to anaphase onset, for RPE1 cells treated with control siRNA, KIF18A siRNA, or KIF18A siRNA + induced expression of WT-GFP-KIF18A. (C) Plot of time in mitosis for the same cells in (B) as a function of polar chromosome presence. Data points represent individual cells and lines indicate median. Data are from three biological replicates and the following number of cells: 69 (control); 77 (KIF18A KD), 74 (KIF18A KD + GFP-KIF18A). Statistical comparisons for B-D were made using Kruskal-Wallis with Dunn’s multiple comparisons test. p-value style: <0.05 (*), <0.01 (**), <0.001 (***), and <0.0001 (****), not significant (>0.05) not shown.

**Figure S2.** The spindle checkpoint protein MAD1 localizes to polar chromosomes in KIF18A KD HeLa cells. (A) Representative images of HeLa cells treated with control siRNA, KIF18A siRNA, or KIF18A siRNA + doxycycline to induce GFP-KIF18A expression (GFP-KIF18A) and labeled with antibodies against CENP-C, MAD1, and GFP. DNA was labeled with DAPI. Scale bar = 10 μm. (B) Plot of the number of kinetochores per HeLa cell that were positive for MAD1. (C) Plot showing the distance between MAD1-positive kinetochores and the metaphase plate in control and KIF18A KD treated cells. (D) Plot of the number of kinetochores per RPE1 cell that were positive for MAD1. Data points represent individual cells and lines indicate median. Data are from three biological replicates and the following number of cells: 44 (control HeLa); 62 (KIF18A KD HeLa), 46 (KIF18A KD + GFP-KIF18A HeLa), 27 (control RPE1); 35 (KIF18A KD RPE1), 60 (KIF18A KD + GFP-KIF18A RPE1). Statistical comparisons for B-D were made using Kruskal-Wallis with Dunn’s multiple comparisons test. p-value style: <0.05 (*), <0.01 (**), <0.001 (***), and <0.0001 (****), not significant (>0.05) not shown.

**Figure S3.** Loss of CENP-E at kinetochores in KIF18A KD cells is associated with mitotic arrest. (A) Representative images of HeLa cells treated with control siRNA, KIF18A siRNA, or KIF18A siRNA + doxycycline to induce GFP-KIF18A then fixed and labeled with antibodies to CENP-C, CENP-E, and GFP. DNA was labeled with DAPI. (B) Plot of CENP-E fluorescence intensity in HeLa cells with the indicated treatment normalized to levels in control siRNA treated cells. Data points represent individual cells and lines indicate median. Data are from three biological replicates and the following number of cells: 47 (control); 55 (KIF18A KD), 47 (KIF18A KD + GFP-KIF18A). Statistical comparisons were made using Kruskal-Wallis with Dunn’s multiple comparisons test. p-value style: <0.05 (*), <0.01 (**), <0.001 (***), and <0.0001 (****), not significant (>0.05) not shown. Scale bars = 10 μm.

**Figure S4.** HT-29 cells continue to proliferate following treatment with 2-4 nM taxol. (A) Plot of normalized cell count over time for HT-29 cells treated with indicated concentrations of Taxol. Lines represent mean and shaded areas represent standard deviation. (B) Plot of fold change in HT-29 cell number from 0 to 96 hours (shown as a % of DMSO control) as a function of Taxol concentration. Data are from three biological replicates. Each dot represents an individual well, N = 18 wells (DMSO) or 9 (all other conditions). Bars indicate mean values. Data were compared by Mann–Whitney test. p-value style: <0.05 (*), <0.01 (**), <0.001 (***), and <0.0001 (****), not significant (>0.05) not shown.

## EXPERIMENTAL MODEL DETAILS

### Cell Lines and Culture Conditions

HT29, SW480, LS1034, HCC1806, HCT116, OVCAR3, RPE1, and MDA-MB-231cells were purchased from ATCC. SKOV3 cells were a generous gift from Alan Howe (University of Vermont). HeLa Kyoto and RPE1 acceptor cells for recombination-mediated cassette exchange (RMCE) were previously described and were a generous gift from Ryoma Ohi (University of Michigan).

All cell lines were authenticated by short tandem repeat (STR) DNA fingerprinting using the Promega GenePrint 10 System according to the manufacturer’s instructions (Promega, #B9510) and were routinely tested for mycoplasma contamination. All cells were cultured at 37 °C in a humidified incubator with 5% CO_2_.

## Culture conditions

- HT29, SW480, and MDA-MB-231 cells were maintained in DMEM/F-12 (Gibco) supplemented with 10% fetal bovine serum (FBS; Gibco) and 1% penicillin-streptomycin.
- HCT116 and SKOV3 cells were cultured in McCoy’s 5A Modified Medium (Gibco) with 10% FBS and 1% penicillin-streptomycin.
- RPE1 and HeLa Kyoto cells were cultured in Minimum Essential Medium Eagle Alpha Modification (MEM-α; Gibco) supplemented with 10% FBS.
- OVCAR3 cells were cultured in RPMI 1640 medium (ATCC) supplemented with 20% FBS, 0.01 mg/mL bovine insulin (Sigma-Aldrich), and 1% penicillin-streptomycin.

## METHOD DETAILS

### siRNA-Mediated Knockdown

siRNA transfections were performed using Lipofectamine RNAiMAX Transfection Reagent (Invitrogen) in Opti-MEM Reduced-Serum Medium (Gibco) following the manufacturer’s protocol. Cells were treated with 5 pmol siRNA per condition.

Cells were treated with specific Silencer Select siRNAs targeting KIF18A (Invitrogen; GCUGGAUUUCAUAAAGUGGtt, CGUUAACUGCAGACGUAAAtt) or a scrambled-sequence negative control siRNA (Invitrogen; Silencer Select Negative Control #1). KIF18A knockdown was previously validated by Western blot or immunofluorescence quantification in all cell types used in this study.

### Small Molecule Inhibitors

Cells were treated with small-molecule inhibitors for 1–2 h prior to fixation and immunostaining unless otherwise noted. Drugs included Sovilnesib (250 nM; Apeiron Therapeutics), MG132 (20 µM; Selleck Chemicals), paclitaxel (Taxol; Selleck Chemicals), reversine (500 nM; Cayman Chemical Company), and nocodazole (5 μm; Selleck Chemicals). For live-cell Sovilnesib spike-in experiments, MG132 was added 10 min prior to imaging, and Sovilnesib was added immediately following the first imaging time point.

### Generation of Inducible KIF18A HeLa and RPE1 Cell Lines Using HILO-RMCE

HeLa Kyoto and RPE1 acceptor cells for high-efficiency, low-background (HILO) recombination-mediated cassette exchange (RMCE) were described previously and maintained in MEM-α with 10% FBS and 10 µg/mL blasticidin. Inducible cell lines expressing siRNA-resistant GFP-KIF18A or GFP-KIF18A^AVVVA^ were generated using Cre-mediated cassette exchange as previously described. GFP-KIF18A^AVVVA^, which disrupts PP1 binding, was generated by mutating residues K612 and W616 to alanine via site-directed mutagenesis, resulting in K612A and W616A, using siRNA-resistant GFP-KIF18A as a template.

Briefly, HeLa Kyoto or RPE1 RMCE acceptor cells were plated at 150,000 cells per well in 6-well plates and co-transfected with a GFP-KIF18A RMCE donor plasmid and a nuclear-localized Cre recombinase plasmid (pEM784) at a 1:10 (wt/wt) ratio using Lipofectamine LTX (Thermo Fisher Scientific). Cells were selected with puromycin (10 µg/mL initially, increased to 20 µg/mL, and then maintained at 5 µg/mL). Correct recombination was confirmed by genomic DNA sequencing. Expression was induced using 2 µg/mL doxycycline for 24 h prior to experiments.

### Kinetochore Microtubule Stability Assays

To assay calcium-stable microtubules, coverslips were collected in room-temperature 1× TBS and immediately transferred to 37 °C calcium chloride buffer for 10 min, followed by fixation in 3.7% paraformaldehyde (Electron Microscopy Sciences) in 1× PHEM buffer for 10 min.

PHEM buffer composition: 60 mM PIPES, 25 mM HEPES, 10 mM EGTA, 4 mM MgSO₄, pH 6.9 (KOH)

Calcium buffer composition: 100 mM PIPES, 1 mM MgCl₂, 1 mM CaCl₂, 0.5% Triton X-100

### Immunofluorescence

Cells were grown on glass coverslips and fixed using one of the following conditions: −20 °C methanol, 1% paraformaldehyde in −20 °C methanol, 2% paraformaldehyde in PHEM, or 4% paraformaldehyde in PHEM. Coverslips were blocked for 1 h with 20% goat serum in antibody dilution buffer (AbDil; TBS, 1% BSA, 0.1% Triton X-100, 0.1% sodium azide).

Primary antibodies included mouse anti-α-tubulin (DM1α; 1:500, Millipore Sigma), guinea pig anti-CENP-C antibody (1:250, Medical and Biological Laboratories), rabbit anti-γ-tubulin (1:500, Abcam), rabbit anti-KIF18A (1:100, Bethyl Laboratories), mouse anti-CENP-E (C-5, Santa Cruz Biotechnology, 1:100,), and mouse anti-MAD1 (9B10, Abcam, 1:400).

Secondary antibodies conjugated to Alexa Fluor 488, 594, and 647 (Molecular Probes) were used at 1:1,000 for 1 h at room temperature. Coverslips were mounted using ProLong Gold antifade mounting medium with DAPI.

### Polar Chromosome Analysis

Cells were seeded onto acid-washed glass coverslips and treated with the indicated siRNAs or drugs for 24 hours prior to fixation. Cells were fixed in 1% paraformaldehyde in −20 °C methanol and stained with mouse anti-γ-tubulin (1:500), rabbit anti-KIF18A (1:100), and guinea pig anti-CENP-C (1:250). Single focal plane images with both spindle poles in focus were acquired. Cells with kinetochores within 1 µm of, or posterior to, spindle poles were scored as positive.

### Chromosome Alignment and Spindle Length Analysis

Cells were treated as described for polar chromosome analysis. Images were acquired with centrosomes in focus. CENP-C fluorescence intensity was quantified along the pole-to-pole axis using ImageJ/Fiji. Intensity profiles were normalized and fitted to Gaussian curves using a custom MATLAB script. Full width at half maximum (FWHM) values for CENP-C fluorescence distribution and spindle length were calculated. Data represent three biological replicates.

### Mitotic Index Analysis

Cells were treated as described for polar chromosome analysis, fixed, and stained. Twenty random fields of view per condition were imaged. Mitotic and total cell numbers were counted manually.

### Proliferation Assays

Cells were plated in 24-well plates and treated with DMSO, sovilnesib (250 nM), taxol, or sovilnesib plus taxol. Plates were imaged every 2 h for 96 h using a BioSpa 8 incubator and Cytation 5 Imaging Reader (BioTek/Agilent). Cell counts were determined using Gen5 software and normalized to control conditions.

### Microtubule Plus-End Tracking

Cells were nucleofected with 2 µg EB3-StayGold plasmid using a 4D-Nucleofector (Lonza). Cells were imaged 24 h later using a Nikon Ti2 spinning-disk microscope at 500 ms intervals for 30 s. EB3 comet tracking and growth velocity measurements were performed using the TrackMate plugin in ImageJ/Fiji.

### Microscopy

Fixed-cell images and live-cell imaging were performed on Ti-2E inverted microscopes (Nikon Instruments) driven by NIS-Elements software (Nikon Instruments). Images were acquired using either a Prime BSI sCMOS camera (Photometrics) or a Zyla sCMOS camera (Andor).

Imaging was conducted using Nikon Plan Apo λ 60X 1.42 NA or APO 100X 1.49 NA oil-immersion objectives for high-resolution fixed-cell imaging. For mitotic index experiments that required lower resolution a Plan Apo 40X 0.95 NA objective was used.

For live-cell imaging experiments, cells were imaged in CO_2_-independent medium (Life Technologies) supplemented with 20% fetal bovine serum (Life Technologies). DNA was labeled with SPY595-DNA (Cytoskeleton). Imaging was performed within environmental chambers maintained at 37 °C.

Long-term live-cell proliferation assays were performed using a Cytation 5 Cell Imaging Multi-Mode Reader (BioTek/Agilent) driven by Gen5 software (BioTek/Agilent). Images were acquired every 2 h using a 4X Plan Fluorite 0.13 NA objective (Olympus) until control cells reached confluency.

### Statistics and Reproducibility

Data are represented as mean ± standard deviation (SD) unless otherwise specified. Statistical tests used to analyze each dataset are described in the figure legends. All statistical tests were performed using GraphPad Prism 10. Exact values for the number of cells/fields analyzed are provided in the figure legends along with the number of independent experimental replicates analyzed.

## References

1. Lengauer, C., Kinzler, K.W., and Vogelstein, B. (1997). Genetic instability in colorectal cancers. Nature 386, 623–627. 10.1038/386623a0.

2. Lee, A.J.X., Endesfelder, D., Rowan, A.J., Walther, A., Birkbak, N.J., Futreal, P.A., Downward, J., Szallasi, Z., Tomlinson, I.P.M., Howell, M., et al. (2011). Chromosomal Instability Confers Intrinsic Multidrug Resistance. Cancer Res 71, 1858–1870. 10.1158/0008-5472.can-10-3604.

3. Bakhoum, S.F., Ngo, B., Laughney, A.M., Cavallo, J.-A., Murphy, C.J., Ly, P., Shah, P., Sriram, R.K., Watkins, T.B.K., Taunk, N.K., et al. (2018). Chromosomal instability drives metastasis through a cytosolic DNA response. Nature 553, 467–472. 10.1038/nature25432.

4. Lengauer, C., Kinzler, K.W., and Vogelstein, B. (1998). Genetic instabilities in human cancers. Nature 396, 643–649. 10.1038/25292.

5. Marquis, C., Fonseca, C.L., Queen, K.A., Wood, L., Vandal, S.E., Malaby, H.L.H., Clayton, J.E., and Stumpff, J. (2021). Chromosomally unstable tumor cells specifically require KIF18A for proliferation. Nat. Commun. 12, 1213. 10.1038/s41467-021-21447-2.

6. Cohen-Sharir, Y., McFarland, J.M., Abdusamad, M., Marquis, C., Bernhard, S.V., Kazachkova, M., Tang, H., Ippolito, M.R., Laue, K., Zerbib, J., et al. (2021). Aneuploidy renders cancer cells vulnerable to mitotic checkpoint inhibition. Nature 590, 486–491. 10.1038/s41586-020-03114-6.

7. Quinton, R.J., DiDomizio, A., Vittoria, M.A., Kotýnková, K., Ticas, C.J., Patel, S., Koga, Y., Vakhshoorzadeh, J., Hermance, N., Kuroda, T.S., et al. (2021). Whole-genome doubling confers unique genetic vulnerabilities on tumour cells. Nature, 1–6. 10.1038/s41586-020-03133-3.

8. Tamayo, N.A., Bourbeau, M.P., Allen, J.R., Ashton, K.S., Chen, J.J., Kaller, M.R., Nguyen, T.T., Nishimura, N., Pettus, L.H., Walton, M., et al. (2022). Targeting the Mitotic Kinesin KIF18A in Chromosomally Unstable Cancers: Hit Optimization Toward an In Vivo Chemical Probe. J. Med. Chem. 65, 4972–4990. 10.1021/acs.jmedchem.1c02030.

9. Payton, M., Belmontes, B., Hanestad, K., Moriguchi, J., Chen, K., McCarter, J.D., Chung, G., Ninniri, M.S., Sun, J., Manoukian, R., et al. (2024). Small-molecule inhibition of kinesin KIF18A reveals a mitotic vulnerability enriched in chromosomally unstable cancers. Nat. Cancer 5, 66–84. 10.1038/s43018-023-00699-5.

10. Schutt, K.L., Queen, K.A., Fisher, K., Budington, O., Mao, W., Liu, W., Gu, X., Xiao, Y., Aswad, F., Joseph, J., et al. (2024). Identification of the KIF18A alpha-4 helix as a therapeutic target for chromosomally unstable tumor cells. Front. Mol. Biosci. 11, 1328077. 10.3389/fmolb.2024.1328077.

11. Sparling, B.A., Lee, H., Zablocki, M.-M., Lynes, M.M., Grigoriu, S., Shehaj, L., Lockbaum, G.J., Khan, S.K., Hotz, T., Lee, Y.-T., et al. (2025). Discovery of Kinesin KIF18A Inhibitor ATX020: Tactical Application of Silicon Atom Replacement. ACS Med. Chem. Lett. 10.1021/acsmedchemlett.5c00512.

12. Phillips, A.F., Zhang, R., Jaffe, M., Schulz, R., Carty, M.C., Verma, A., Feinberg, T.Y., Arensman, M.D., Chiu, A., Letso, R., et al. (2025). Targeting chromosomally unstable tumors with a selective KIF18A inhibitor. Nat. Commun. 16, 307. 10.1038/s41467-024-55300-z.

13. Stumpff, J., du, Y., English, C.A., Maliga, Z., Wagenbach, M., Asbury, C.L., Wordeman, L., and ohi, R. (2011). A tethering mechanism controls the processivity and kinetochore-microtubule plus-end enrichment of the kinesin-8 Kif18A. Mol Cell 43, 764–775. 10.1016/j.molcel.2011.07.022.

14. Stumpff, J., Dassow, G. von, Wagenbach, M., Asbury, C., and Wordeman, L. (2008). The Kinesin-8 Motor Kif18A Suppresses Kinetochore Movements to Control Mitotic Chromosome Alignment. Dev. Cell 14, 252–262. 10.1016/j.devcel.2007.11.014.

15. Stumpff, J., Wagenbach, M., Franck, A., Asbury, C.L., and Wordeman, L. (2012). Kif18A and Chromokinesins Confine Centromere Movements via Microtubule Growth Suppression and Spatial Control of Kinetochore Tension. Dev. Cell 22, 1017–1029. 10.1016/j.devcel.2012.02.013.

16. Fonseca, C.L., Malaby, H.L.H., Sepaniac, L.A., Martin, W., Byers, C., Czechanski, A., Messinger, D., Tang, M., Ohi, R., Reinholdt, L.G., et al. (2019). Mitotic chromosome alignment ensures mitotic fidelity by promoting interchromosomal compaction during anaphase. J. Cell Biol. 218, 1148–1163. 10.1083/jcb.201807228.

17. Mayr, M.I., Hümmer, S., Bormann, J., Grüner, T., Adio, S., Woehlke, G., and Mayer, T.U. (2007). The Human Kinesin Kif18A Is a Motile Microtubule Depolymerase Essential for Chromosome Congression. Curr. Biol. 17, 488–498. 10.1016/j.cub.2007.02.036.

18. Czechanski, A., Kim, H., Byers, C., Greenstein, I., Stumpff, J., and Reinholdt, L.G. (2015). Kif18a is specifically required for mitotic progression during germ line development. Dev Biol 402, 253–262. 10.1016/j.ydbio.2015.03.011.

19. King, R.W., Peters, J.-M., Tugendreich, S., Rolfe, M., Hieter, P., and Kirschner, M.W. (1995). A 20s complex containing CDC27 and CDC16 catalyzes the mitosis-specific conjugation of ubiquitin to cyclin B. Cell 81, 279–288. 10.1016/0092-8674(95)90338-0.

20. Sudakin, V., Ganoth, D., Dahan, A., Heller, H., Hershko, J., Luca, F.C., Ruderman, J.V., and Hershko, A. (1995). The cyclosome, a large complex containing cyclin-selective ubiquitin ligase activity, targets cyclins for destruction at the end of mitosis. Mol. Biol. Cell 6, 185–197. 10.1091/mbc.6.2.185.

21. Sudakin, V., Chan, G.K.T., and Yen, T.J. (2001). Checkpoint inhibition of the APC/C in HeLa cells is mediated by a complex of BUBR1, BUB3, CDC20, and MAD2. J. Cell Biol. 154, 925–936. 10.1083/jcb.200102093.

22. Li, X., and Nicklas, R.B. (1995). Mitotic forces control a cell-cycle checkpoint. Nature 373, 630–632. 10.1038/373630a0.

23. Nesbit, C., Martin, W., Czechanski, A., Byers, C., Raghupathy, N., Ferraj, A., Stumpff, J., and Reinholdt, L.G. (2024). Anapc5andAnapc7as genetic modifiers of KIF18A function in fertility and mitotic progression. 10.1101/2024.12.03.626395.

24. Gliech, C.R., Yeow, Z.Y., Tapias-Gomez, D., Yang, Y., Huang, Z., Tijhuis, A.E., Spierings, D.C., Foijer, F., Chung, G., Tamayo, N., et al. (2024). Weakened APC/C activity at mitotic exit drives cancer vulnerability to KIF18A inhibition. EMBO J., 1–29. 10.1038/s44318-024-00031-6.

25. Queen, K.A., Cario, A., Berger, C.L., and Stumpff, J. (2024). Modification of the neck-linker of KIF18A alters Microtubule subpopulation preference. Mol. Biol. Cell 35, ar3. 10.1091/mbc.e23-05-0167.

26. Vukušić, K., and Tolić, I.M. (2022). Polar Chromosomes—Challenges of a Risky Path. Cells 11, 1531. 10.3390/cells11091531.

27. Mitchison, T., Evans, L., Schulze, E., and Kirschner, M. (1986). Sites of microtubule assembly and disassembly in the mitotic spindle. Cell 45, 515–527. 10.1016/0092-8674(86)90283-7.

28. Kapoor, T.M., Mayer, T.U., Coughlin, M.L., and Mitchison, T.J. (2000). Probing Spindle Assembly Mechanisms with Monastrol, a Small Molecule Inhibitor of the Mitotic Kinesin, Eg5. J. Cell Biol. 150, 975–988. 10.1083/jcb.150.5.975.

29. DeLuca, J.G., Dong, Y., Hergert, P., Strauss, J., Hickey, J.M., Salmon, E.D., and McEwen, B.F. (2005). Hec1 and nuf2 are core components of the kinetochore outer plate essential for organizing microtubule attachment sites. Mol Biol Cell 16, 519–531. 10.1091/mbc.e04-09-0852.

30. DeLuca, J.G., Gall, W.E., Ciferri, C., Cimini, D., Musacchio, A., and Salmon, E.D. (2006). Kinetochore microtubule dynamics and attachment stability are regulated by Hec1. Cell 127, 969–982. 10.1016/j.cell.2006.09.047.

31. Cheeseman, I.M., Chappie, J.S., Wilson-Kubalek, E.M., and Desai, A.B. (2006). The conserved KMN network constitutes the core microtubule-binding site of the kinetochore. Cell 127, 983–997. 10.1016/j.cell.2006.09.039.

32. DeLuca, K.F., Lens, S.M.A., and DeLuca, J.G. (2011). Temporal changes in Hec1 phosphorylation control kinetochore–microtubule attachment stability during mitosis. J Cell Sci 124, 622–634. 10.1242/jcs.072629.

33. Häfner, J., Mayr, M.I., Möckel, M.M., and Mayer, T.U. (2014). Pre-anaphase chromosome oscillations are regulated by the antagonistic activities of Cdk1 and PP1 on Kif18A. Nat Commun 5, 4397. 10.1038/ncomms5397.

34. Meadows, J.C., Shepperd, L.A., Vanoosthuyse, V., Lancaster, T.C., Sochaj, A.M., Buttrick, G.J., Hardwick, K.G., and Millar, J.B.A. (2011). Spindle checkpoint silencing requires association of PP1 to both Spc7 and kinesin-8 motors. Dev Cell 20, 739–750. 10.1016/j.devcel.2011.05.008.

35. Huang, Y., Yao, Y., Xu, H.-Z., Wang, Z.-G., Lu, L., and Dai, W. (2009). Defects in chromosome congression and mitotic progression in KIF18A-deficient cells are partly mediated through impaired functions of CENP-E. Cell cycle (Georgetown, Tex.) 8, 2643–2649.

36. Liu, X. s, Zhao, X. d, Wang, X., Yao, Y. x, Zhang, L. l, Shu, R. z, Ren, W. h, Huang, Y., Huang, L., Gu, M. m, et al. (2010). Germinal Cell Aplasia in Kif18a Mutant Male Mice Due to Impaired Chromosome Congression and Dysregulated BubR1 and CENP-E. Genes Cancer 1, 26–39. 10.1177/1947601909358184.

37. Miao, J., Hara, M., Su, K.-C., Keys, H.R., Kong, W., Takenoshita, Y., Cheeseman, I.M., and Fukagawa, T. (2025). KIF18A promotes chromosome congression in cooperation with CENP-E downstream of CENP-C. Cell Rep., 116515. 10.1016/j.celrep.2025.116515.

38. Wood, K.W., Sakowicz, R., Goldstein, L.S., and Cleveland, D.W. (1997). CENP-E is a plus end-directed kinetochore motor required for metaphase chromosome alignment. Cell 91, 357–366.

39. Schaar, B.T., Chan, G.K., Maddox, P., Salmon, E.D., and Yen, T.J. (1997). CENP-E function at kinetochores is essential for chromosome alignment. The Journal of Cell Biology 139, 1373–1382.

40. Kapoor, T.M., Lampson, M.A., Hergert, P., Cameron, L., Cimini, D., Salmon, E.D., McEwen, B.F., and Khodjakov, A. (2006). Chromosomes can congress to the metaphase plate before biorientation. Science 311, 388–391. 10.1126/science.1122142.

41. Gudimchuk, N., Vitre, B., Kim, Y., Kiyatkin, A., Cleveland, D.W., Ataullakhanov, F.I., and Grishchuk, E.L. (2013). Kinetochore kinesin CENP-E is a processive bi-directional tracker of dynamic microtubule tips. Nat Cell Biol 15. 10.1038/ncb2831.

42. Yen, T.J., Li, G., Schaar, B.T., Szilak, I., and Cleveland, D.W. (1992). CENP-E is a putative kinetochore motor that accumulates just before mitosis. Nature 359, 536–539. 10.1038/359536a0.

43. Howell, B.J., McEwen, B.F., Canman, J.C., Hoffman, D.B., Farrar, E.M., Rieder, C.L., and Salmon, E.D. (2001). Cytoplasmic dynein/dynactin drives kinetochore protein transport to the spindle poles and has a role in mitotic spindle checkpoint inactivation. J Cell Biology 155, 1159–1172. 10.1083/jcb.200105093.

44. Ertych, N., Stolz, A., Stenzinger, A., Weichert, W., Kaulfuß, S., Burfeind, P., Aigner, A., Wordeman, L., and Bastians, H. (2014). Increased microtubule assembly rates influence chromosomal instability in colorectal cancer cells. Nat. Cell Biol. 16, 779–791. 10.1038/ncb2994.

45. du, Y., English, C.A., and ohi, R. (2010). The kinesin-8 Kif18A dampens microtubule plus-end dynamics. Curr Biol 20, 374–380. 10.1016/j.cub.2009.12.049.

46. Armond, J.W., Vladimirou, E., Erent, M., McAinsh, A.D., and Burroughs, N.J. (2015). Probing microtubule polymerisation state at single kinetochores during metaphase chromosome motion. J Cell Sci 128, 1991–2001. 10.1242/jcs.168682.

47. Wordeman, L., Decarreau, J., Vicente, J.J., and Wagenbach, M. (2016). Divergent microtubule assembly rates after short-versus long-term loss of end-modulating kinesins. Mol Biol Cell 27, 1300–1309. 10.1091/mbc.e15-11-0803.

48. Fuller, B.G., Lampson, M.A., Foley, E.A., Rosasco-Nitcher, S., Le, K.V., Tobelmann, P., Brautigan, D.L., Stukenberg, P.T., and Kapoor, T.M. (2008). Midzone activation of aurora B in anaphase produces an intracellular phosphorylation gradient. Nature 453, 1132–1136. 10.1038/nature06923.

49. Kuhn, J., and Dumont, S. (2019). Mammalian kinetochores count attached microtubules in a sensitive and switch-like manner. The Journal of Cell Biology 218, 3583–3596. 10.1083/jcb.201902105.

