## Supplemental Figures for "KIF18A Maintains Kinetochore-Microtubule Attachments in CIN Cells by Limiting Microtubule Polymerization"

Figure S1.

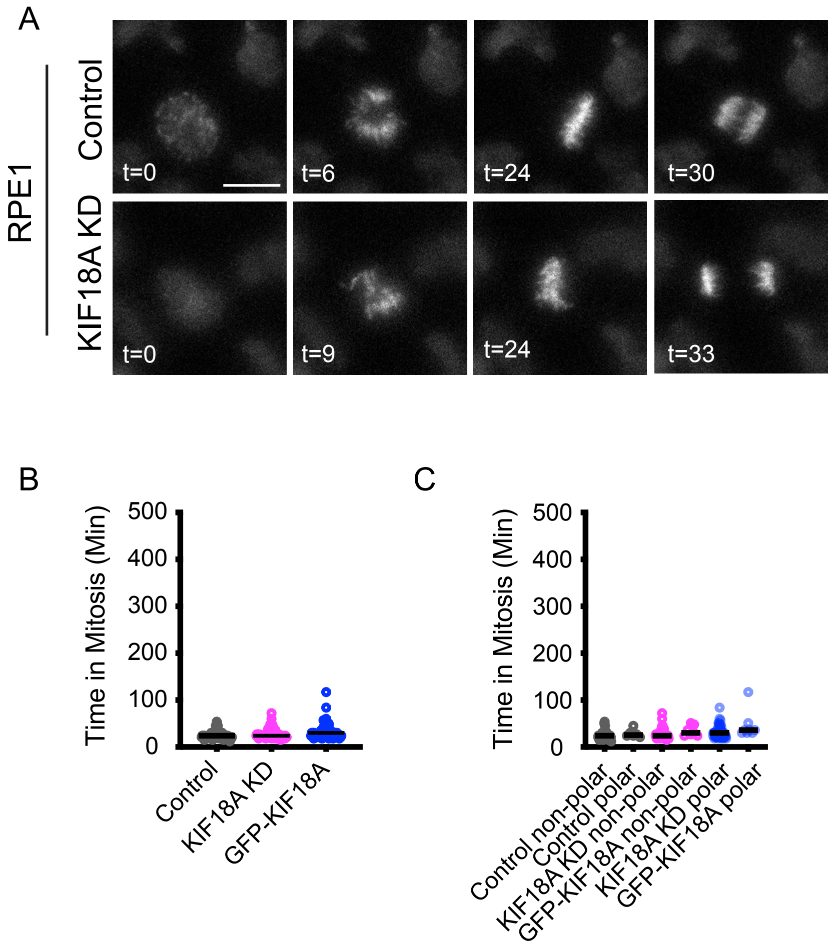
A
RPE1
Control
KIF18A KD
t=0
t=6
t=24
t=30
t=0
t=9
t=24
t=33
B
Time in Mitosis (Min)
500
400
300
200
100
0
Control
KIF18A KD
GFP-KIF18A
C
Time in Mitosis (Min)
500
400
300
200
100
0
Control non-polar
Control polar
KIF18A KD non-polar
GFP-KIF18A non-polar
KIF18A KD polar
GFP-KIF18A polar

**Figure S1. KIF18A KD does not induce polar chromosomes in RPE1 cells**. (A) Representative still frames from time-lapse imaging of chromosomes in RPE1 cells labeled with SPY-DNA. Time (t) in minutes relative to chromosome condensation is indicated. Scale bar = 10 μm. (B) Plot of time in mitosis, measured from chromosome condensation to anaphase onset, for RPE1 cells treated with control siRNA, KIF18A siRNA, or KIF18A siRNA + induced expression of WT-GFP-KIF18A. (C) Plot of time in mitosis for the same cells in (B) as a function of polar chromosome presence. Data points represent individual cells and lines indicate median. Data are from three biological replicates and the following number of cells: 69 (control); 77 (KIF18A KD), 74 (KIF18A KD + GFP-KIF18A). Statistical comparisons for B-D were made using Kruskal-Wallis with Dunn's multiple comparisons test. p-value style: <0.05 (*), <0.01 (**), <0.001 (***), and <0.0001 (****), not significant (>0.05) not shown.

Figure S2.

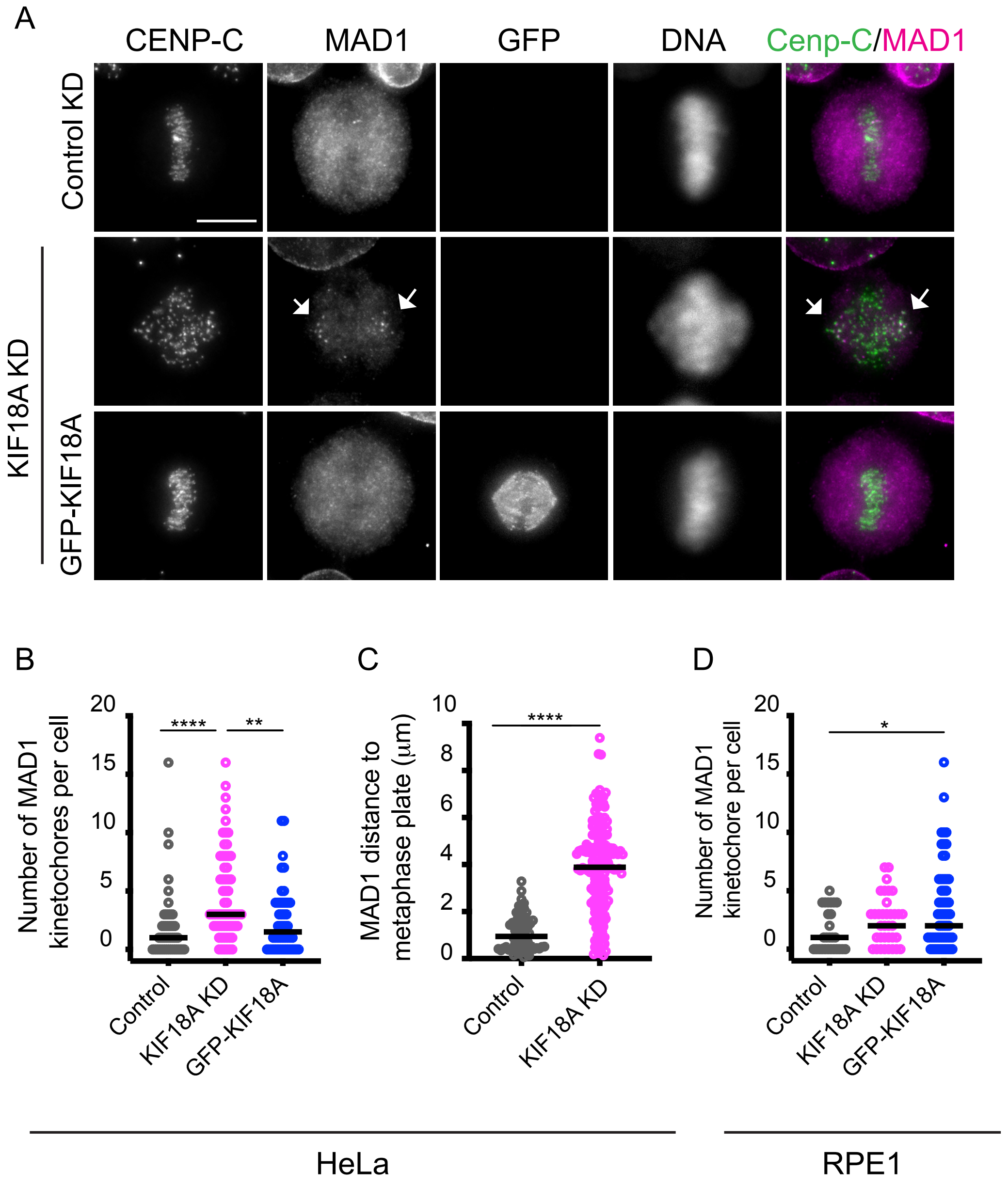
A
CENP-C
MAD1
GFP
DNA
Cenp-C/MAD1
Control KD
KIF18A KD
GFP-KIF18A
B
Number of MAD1 kinetochores per cell
20
15
10
5
0
****
**
Control
KIF18A KD
GFP-KIF18A
C
MAD1 distance to metaphase plate (μm)
10
8
6
4
2
0
****
Control
KIF18A KD
D
Number of MAD1 kinetochore per cell
20
15
10
5
0
*
Control
KIF18A KD
GFP-KIF18A
HeLa
RPE1

**Figure S2. The spindle checkpoint protein MAD1 localizes to polar chromosomes in KIF18A KD HeLa cells.** (A) Representative images of HeLa cells treated with control siRNA, KIF18A siRNA, or KIF18A siRNA + doxycycline to induce GFP-KIF18A expression (GFP-KIF18A) and labeled with antibodies against CENP-C, MAD1, and GFP. DNA was labeled with DAPI. Scale bar = 10 μm. (B) Plot of the number of kinetochores per HeLa cell that were positive for MAD1. (C) Plot showing the distance between MAD1-positive kinetochores and the metaphase plate in control and KIF18A KD treated cells. (D) Plot of the number of kinetochores per RPE1 cell that were positive for MAD1. Data points represent individual cells and lines indicate median. Data are from three biological replicates and the following number of cells: 44 (control HeLa); 62 (KIF18A KD HeLa), 46 (KIF18A KD + GFP-KIF18A HeLa), 27 (control RPE1); 35 (KIF18A KD RPE1), 60 (KIF18A KD + GFP-KIF18A RPE1). Statistical comparisons for B-D were made using Kruskal-Wallis with Dunn's multiple comparisons test. p-value style: $<0.05$ (*), $<0.01$ (**), $<0.001$ (***), and $<0.0001$ (****), not significant ($>0.05$) not shown.

Figure S3.

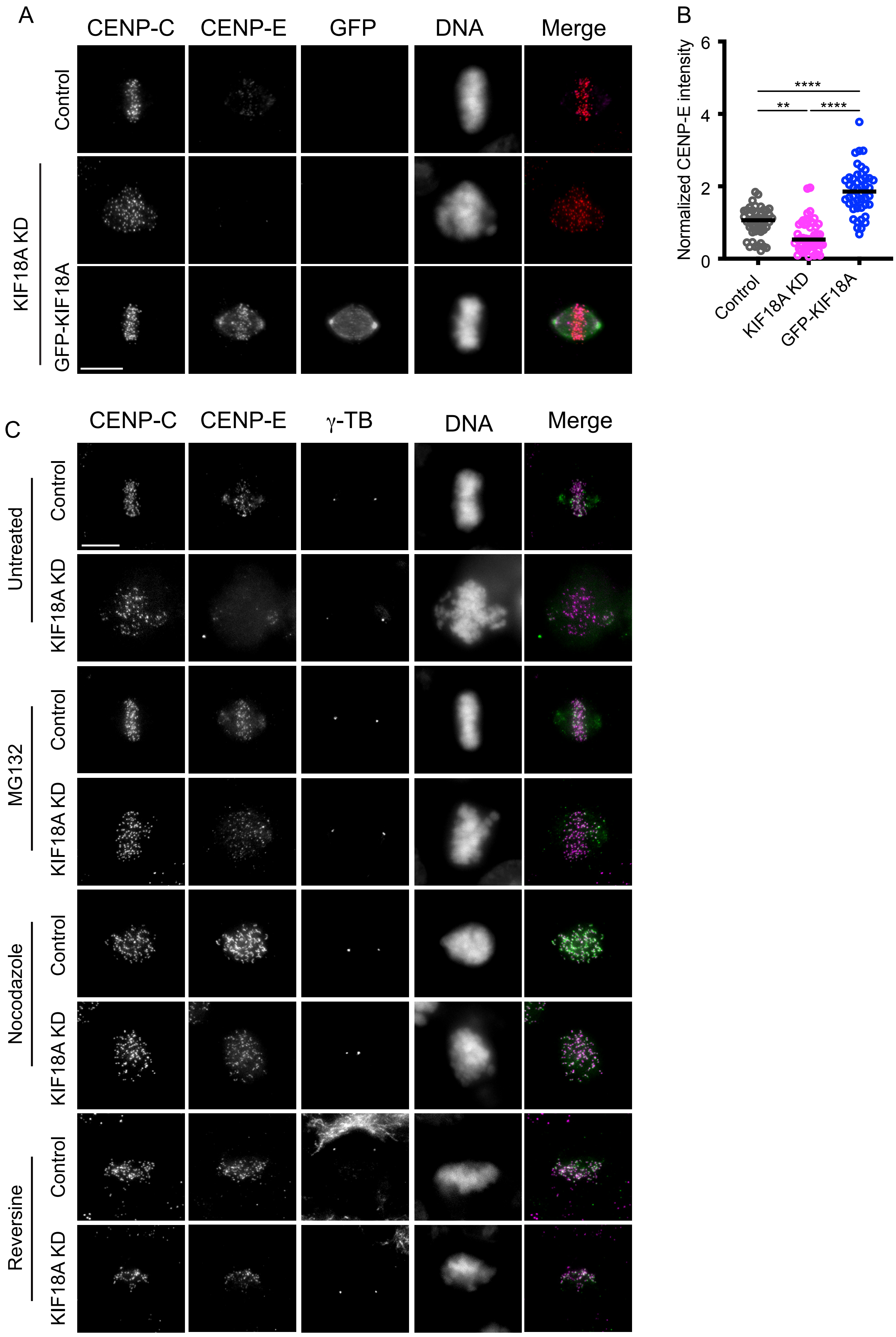
A
CENP-C
CENP-E
GFP
DNA
Merge
Control
KIF18A KD
GFP-KIF18A
B
Normalized CENP-E intensity
6
4
2
0
****
**
****
Control
KIF18A KD
GFP-KIF18A
C
CENP-C
CENP-E
γ-TB
DNA
Merge
Untreated
Control
KIF18A KD
MG132
Control
KIF18A KD
Nocodazole
Control
KIF18A KD
Reversine
Control
KIF18A KD

**Figure S3. Loss of CENP-E at kinetochores in KIF18A KD cells is associated with mitotic arrest.** (A) Representative images of HeLa cells treated with control siRNA, KIF18A siRNA, or KIF18A siRNA + doxycycline to induce GFP-KIF18A then fixed and labeled with antibodies to CENP-C, CENP-E, and GFP. DNA was labeled with DAPI. (B) Plot of CENP-E fluorescence intensity in HeLa cells with the indicated treatment normalized to levels in control siRNA treated cells. Data points represent individual cells and lines indicate median. Data are from three biological replicates and the following number of cells: 47 (control); 55 (KIF18A KD), 47 (KIF18A KD + GFP-KIF18A). Statistical comparisons were made using Kruskal-Wallis with Dunn's multiple comparisons test. p-value style: <0.05 (*), <0.01 (**), <0.001 (***), and <0.0001 (****), not significant (>0.05) not shown. Scale bars = 10 μm.

Figure S4.

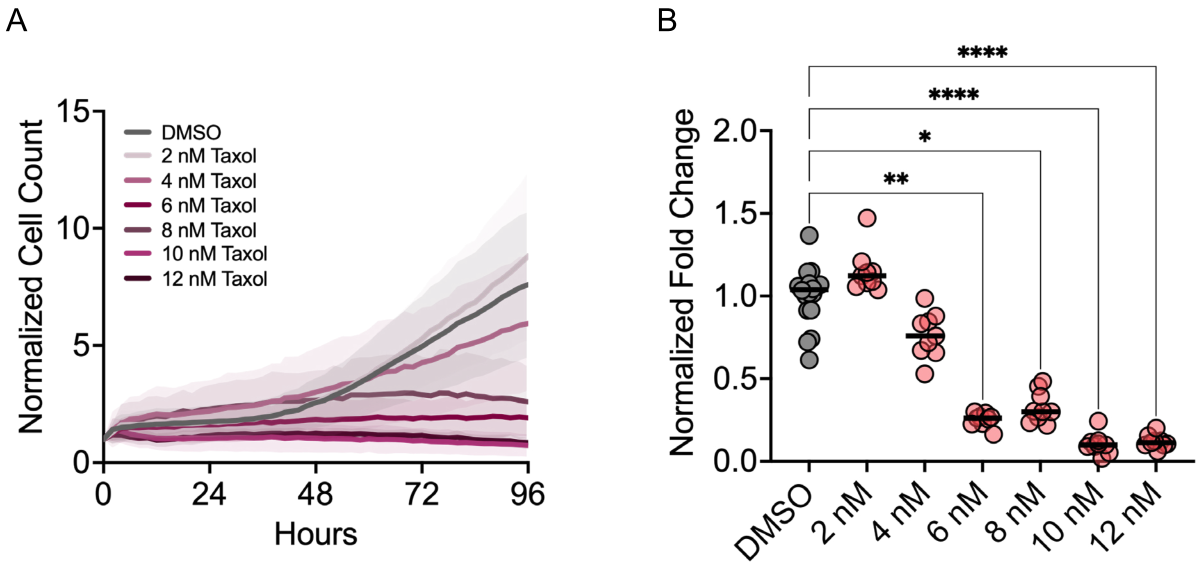

A
15
10
5
0
Normalized Cell Count
DMSO
2 nM Taxol
4 nM Taxol
6 nM Taxol
8 nM Taxol
10 nM Taxol
12 nM Taxol
0
24
48
72
96
Hours
B
****
****
*
**
2.0
1.5
1.0
0.5
0.0
Normalized Fold Change
DMSO
2 nM
4 nM
6 nM
8 nM
10 nM
12 nM

**Figure S4. HT-29 cells continue to proliferate following treatment with 2-4 nM taxol**. (A) Plot of normalized cell count over time for HT-29 cells treated with indicated concentrations of Taxol. Lines represent mean and shaded areas represent standard deviation. (B) Plot of fold change in HT-29 cell number from 0 to 96 hours (shown as a % of DMSO control) as a function of Taxol concentration. Data are from three biological replicates. Each dot represents an individual well, N = 18 wells (DMSO) or 9 (all other conditions). Bars indicate mean values. Data were compared by Mann–Whitney test. p-value style: <0.05 (*), <0.01 (**), <0.001 (***), and <0.0001 (****), not significant (>0.05) not shown.
